# Cholinergic Modulation of Hippocampally Mediated Attention and Perception

**DOI:** 10.1101/2020.05.19.104497

**Authors:** Nicholas A. Ruiz, Monica Thieu, Mariam Aly

**Affiliations:** Department of Psychology, Columbia University; Affiliate Member, Zuckerman Mind Brain Behavior Institute, Columbia University

**Keywords:** Relational attention, selective attention, encoding, neuromodulation, medial temporal lobe

## Abstract

Attention to the relations between visual features modulates hippocampal representations. Moreover, hippocampal damage impairs discrimination of spatial relations. We explore a mechanism by which this might occur: modulation by the acetylcholine system. Acetylcholine enhances afferent input to the hippocampus and suppresses recurrent connections within it. This biases hippocampal processing toward environmental input, and should improve externally-oriented, hippocampally mediated attention and perception. We examined cholinergic modulation on an attention task that recruits the hippocampus. On each trial, participants viewed two images (rooms with paintings). On “similar room” trials, they judged whether the rooms had the same spatial layout from a different perspective. On “similar art” trials, they judged whether the paintings could have been painted by the same artist. On “identical” trials, participants simply had to detect identical paintings or rooms. We hypothesized that cholinergic modulation would improve performance on the similar room task, given past findings that hippocampal representations predicted, and hippocampal damage impaired, behavior on this task. To test this, nicotine cigarette smokers took part in two sessions: one before which they abstained from nicotine for 12 hours, and one before which they ingested nicotine in the past hour. Individual differences in expired breath carbon monoxide levels — a measure of how recently or how much someone smoked — predicted performance improvements on the similar room task. This finding provides novel support for computational models that propose that acetylcholine enhances externally oriented attentional states in the hippocampus.

## Introduction

Imagine that you are in an art museum. You might choose to direct your attention to a specific painting on the wall. However, once you do that, the visual features of that painting might trigger memory retrieval of associated information — e.g., similar paintings that you have seen in other museums. In order to better focus your attention on the painting in front of you, you may wish to suppress memories of other related paintings. How do we balance the need to attend to the external world vs our internal memories?

Many regions of the brain balance this need by switching between internal vs external processing modes (Honey, Newman, & Schapiro, 2017). For example, the hippocampus oscillates between states that are optimized for attention to the external world (and the encoding of new memories) and states that are optimized for memory retrieval (Decker & Duncan, 2020; <u>Hasselmo, Bodelón, & Wyble, 2002</u>; Tarder-Stoll, Jayakumar, Dimsdale-Zucker, Günseli, & Aly, 2020). These distinct attention/encoding and memory retrieval modes are thought to be coordinated, at least in part, by the acetylcholine neurotransmitter system. Evidence from computational models and electrophysiology studies in rodents have shown that high levels of acetylcholine bias the hippocampus toward attention/encoding, and low levels of acetylcholine bias the hippocampus toward memory retrieval (**Figure 1;** Easton, Douchamps, Eacott, & Lever 2012; Hasselmo & Schnell 1994; Hasselmo, 1995; Hasselmo & Barkai, 1995; Hasselmo, Schnell, & Barkai, 1995; Hasselmo, Wyble, & Wallenstein, 1996; Hasselmo & McGaughy, 2004; Meeter, Murre, & Talamini, 2004; Newman, Gupta, Climer, Monaghan, & Hasselmo, 2012).

**Figure 1.**
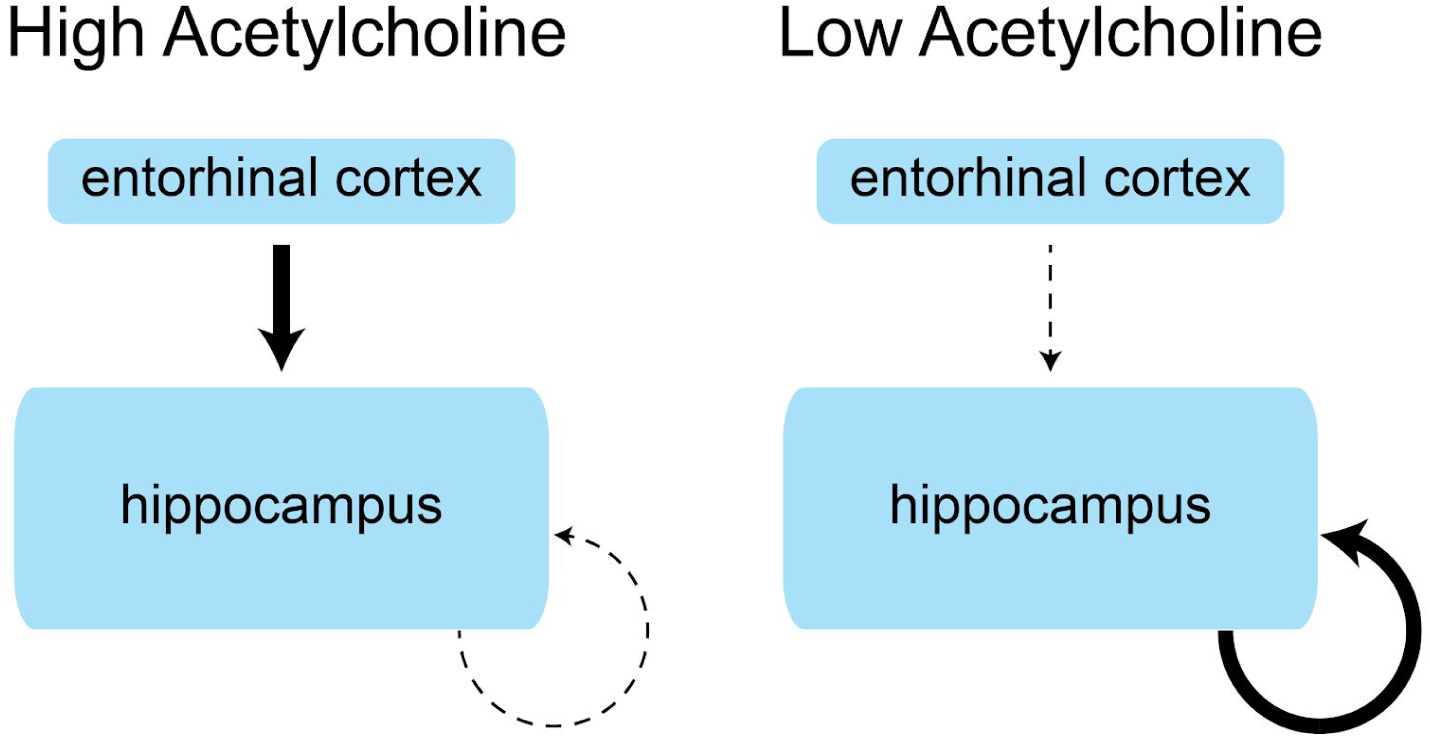
Cholinergic modulation of hippocampal function. High levels of acetylcholine (left) strengthen afferent input from entorhinal cortex to the hippocampus (thick arrow) — a flow of input important for externally oriented attention and encoding. This is accompanied by the suppression of recurrent connections within the hippocampus (particularly subfield CA3; dashed arrow) — an excitatory circuit important for memory retrieval and internally oriented processing. The result of these two mechanisms biases the hippocampus towards a state that prioritizes attention and encoding. Conversely, low levels of acetylcholine (right) prioritize recurrent connections within the hippocampus and suppress input from entorhinal cortex. This biases the hippocampus towards a retrieval state. This figure is adapted from Hasselmo (2006) and Newman et al., (2012).

In particular, high levels of acetylcholine prioritize afferent input from entorhinal cortex to the hippocampus, and lower the threshold for long-term potentiation (LTP) in both entorhinal cortex (Yun et al., 2000) and hippocampal subfield CA1 (Huerta & Lisman, 1993). Additionally, high levels of acetylcholine suppress excitatory recurrent connections in hippocampal subfield CA3 (Hasselmo et al., 1995). Together, these changes should enhance attention to, and perception of, the external world (and encoding of new memories): sensory signals received and integrated by entorhinal cortex (<u>Lavanex & Amaral, 2000</u>) are given prioritized processing, while pattern completion mechanisms in CA3 are suppressed (Hasselmo et al., 1995), impairing memory retrieval (also see Kukolja, Thiel, & Fink, 2009).

Conversely, low levels of acetylcholine prioritize excitatory recurrent connections in CA3 and suppress afferent input from entorhinal cortex to the hippocampus (Figure 1; Hasselmo & Schnell 1994; Newman et al., 2012). This biases the hippocampus toward a retrieval state.

This framework has been supported by studies in rodents that employ the use of cholinergic agonists and antagonists, as well as by computational modeling (Hasselmo et al., 1995; Hasselmo & Schnell, 1994). However, due to the difficulty of conducting pharmacological manipulations, these models have rarely been directly tested in humans, e.g., with direct manipulations of the acetylcholine system. One notable study that tested this model found that scopolamine, an antagonist of the muscarinic acetylcholine receptor, impaired new learning and increased proactive interference, while not affecting retrieval of previously learned information (Atri et al., 2004). This study therefore offers evidence that reduced functioning of the acetylcholine system can impair attention/encoding but does not affect memory retrieval. The goal of the current study is to investigate a complementary issue: whether cholinergic enhancement can improve hippocampally mediated attention and perception.

Although few studies in humans have directly tested cholinergic modulation of hippocampal attention/encoding vs retrieval states, some research has indirectly tested these models by using behavior as a window into cholinergic modulation. Specifically, these studies have taken advantage of the fact that the effects of cholinergic modulation are extended in time (Hasselmo & Fehlau, 2001) and more acetylcholine is released in the hippocampus when rodents explore a novel vs familiar environment (Giovannini et al., 2001). Thus, exposure to novel stimuli should increase acetylcholine release and bias the hippocampus toward an encoding state. Conversely, exposure to familiar stimuli should be associated with a relative decrease in acetylcholine release, biasing the hippocampus toward a retrieval state. Studies relying on this logic have found that recent exposure to novel stimuli improves behaviors that depend on encoding precise memories (Duncan, Sadanand, & Davachi, 2012), while recent exposure to familiar stimuli improves behaviors that depend on memory retrieval (Duncan et al., 2012; Duncan & Shohamy, 2016; <u>Patil & Duncan, 2018</u>). These studies are consistent with cholinergic modulation models of hippocampal encoding and retrieval states (Hasselmo et al., 1995; Hasselmo & Schnell 1994), but they relied on indirect manipulations of acetylcholine (via novelty).

Motivated by these two classes of studies, here we directly test the prediction that cholinergic agonists should enhance hippocampally mediated attention and perception, in a task with no demands on long-term memory. We capitalized on an attention task that we have previously shown recruits the hippocampus (Aly & Turk-Browne, 2016a; Aly & Turk-Browne, 2016b; Ruiz, Meager, Agarwal, & Aly, 2020). On each trial, participants view two 3D-rendered rooms, each with distinct shapes, different pieces of furniture, and a single painting. Participants are required to attend either to the spatial features of the rooms (i.e., angles of the walls, arrangement of the furniture; “room” trials) or to the artistic style of the paintings (i.e., use of color, brushstrokes, amount of detail; “art” trials). Within these two attention tasks, there were two trial types that varied in difficulty and complexity. On “similar” trials, individuals were required to identify non-identical but similar rooms or paintings. Specifically, on similar room trials, participants had to identify rooms with the same spatial layout from a different perspective, although other visual features (wall color, furniture exemplars) were altered. On similar art trials, participants had to identify different paintings that were painted by the same artist. These paintings had similar style (use of color, type of brushstroke), but their content could differ. On “identical” trials, participants simply needed to detect identical repetitions of a room (identical room trials) or identical repetitions of a painting (identical art trials).

We have previously found that the stability of hippocampal activity patterns across similar room trials predicts performance on those trials, but no such relationship exists for similar art trials (Aly & Turk-Browne, 2016a; Aly & Turk-Browne, 2016b). Furthermore, performance on similar room trials, but not similar art trials or identical trials, is dependent on an intact hippocampus/medial temporal lobe (Ruiz et al., 2020). This may be because the similar room task places higher demands on relational representations than the other tasks, and relational processing is a critical aspect of hippocampal function (<u>Aly & Turk-Browne, 2018</u>; Eichenbaum & Cohen, 2014; Konkel & Cohen, 2009; Olsen, Moses, Riggs, & Ryan, 2012). We note here that, although this is an attention task that requires externally oriented processing, it may nevertheless benefit from some amount of internally oriented processing as well. We will return to this important point in the Discussion.

If cholinergic modulation enhances hippocampal attention and perception, then individuals should perform better on the similar room task when they have ingested a cholinergic agonist, compared to when they have not. To test this, we examined how performance on these trial types is affected by nicotine, a cholinergic agonist (Brody et al., 2006) that has been linked to hippocampal learning and memory (Hasselmo, 2006; Kutlu and Gould, 2015; Levin, Addy, & Sigurani, 2002; Newman et al., 2012; Ohno, Yamamoto, & Wantanabe, 1993; Placzek, Zhang, & Dani, 2009).

To that end, we conducted a study with nicotine cigarette smokers. We chose this population because it provides a relatively tractable way of conducting pharmacological manipulations in humans (note that another tractable approach would be to manipulate caffeine intake, because caffeine can increase acetylcholine levels; Carter, O’Connor, Carter, & Ungerstedt, 1995). Nicotine cigarette smokers came into our lab for 2 sessions, one before which they had just smoked a cigarette (“ON” session) and another before which they had abstained from smoking for at least 12 hours (“OFF” session). Upon arrival, participants were tested for compliance via an expired breath carbon monoxide (CO) monitor. CO levels rise with tobacco smoking (Jarvis, Russell, & Saloojee, 1980; Wald, Idle, Boreham, & Bailey, 1981), thus providing a measure of how recently or how much an individual has smoked and an index of nicotine levels (Jarvik et al., 2000; Russell, Martin, Taylor, Feyerabend, & Saloojee, 1978; Vollstädt-Klein et al., 2011; Vossel, Warbrick, Mobascher, Winterer, & Fink, 2011). Participants then filled out a short questionnaire inquiring how many cigarettes they smoked in the last hour and the last 12 hours. Finally, they completed the attention task described above (also see Ruiz et al., 2020).

Our main prediction was that performance on the (hippocampally mediated) similar room task should improve when individuals are on vs. off nicotine. A complementary hypothesis is that performance enhancements on similar room trials might scale with the amount of nicotine ingested. If so, the recency or amount of smoking, as indexed by expired breath CO levels, might predict performance improvements. Both of these findings would be consistent with cholinergic modulation of hippocampal attention and/or perception. An alternative hypothesis is that nicotine will enhance performance in a non-selective manner across all trial types, given findings that cholinergic signaling can modulate visual cortex responses, e.g., via gain modulation or sharpening (Disney, Aoki, & Hawken, 2007; Sarter, Hasselmo, Bruno, & Givens, 2005; Silver, Shenhav, & D’Esposito, 2008) and improve visual attention and perception (Ernst, Heishman, Spurgeon, & London, 2001; Gratton et al., 2017; Hahn et al., 2007; Parrott & Roberts, 1991).

## Materials and Methods

### Participants

Nicotine cigarette smokers (n = 60) were recruited via flyers posted around the Columbia University community. The study was approved by the Columbia University Institutional Review Board, and participants received monetary compensation for their time. Inclusion criteria were: smoking at least one cigarette per week; at least 18 years of age; normal or corrected-to-normal vision; and fluency in English. The exclusion criteria were (1) smoking e-cigarettes, because such cigarettes do not leave carbon monoxide (CO) traces in expired breath — a measure we used to assess compliance; and (2) having an expired breath CO level during the OFF session that was higher than or equal to the CO level during the ON session (see **Procedure**). Ten participants were excluded for this reason. Data for the remaining fifty participants are reported here (19 women, 31 men; *M*_*age*_ = 25.7 years; *M*_*education*_ = 16.3 years). Participants ranged in how many cigarettes they smoked per day, from less than 1 to 14 (*M* = 4.60, *SD* = 3.67) and ranged in how long they have been smoking, from 4 months to 20 years (*M* = 6.75 years, *SD =* 4.95). (Here and elsewhere, *M* refers to the mean and *SD* to the standard deviation).

### Questionnaires

Participants completed the Fagerstrom Test for Nicotine Dependence (FTND; Heatherton, Kozlowski, Frecker, & Fagerstrom, 1991). FTND questions assess: how soon the individual smokes their first cigarette upon waking up; if the individual has difficulty giving up cigarettes in places where smoking is forbidden (e.g., library, movie theater); if the individual would rather give up the first cigarette of the day or any other; how many cigarettes the individual smokes per day; if the individual smokes more in the morning or during the rest of the day; and if the individual smokes when they are ill. Yes/no items are scored as 0 (no) or 1 (yes), and multiple-choice items are scored from 0-3. All participants scored in the “low dependence” or “low to moderate dependence” range on this assessment (*M* = 1.02, *SD* = 1.11; 1-2 = low dependence, 3-4 = low to moderate dependence, 5-7 = moderate dependence, 8-10 = high dependence). Because no participant scored in the moderate or high dependence range, we believe it is unlikely that severe withdrawal symptoms occurred in the OFF nicotine session Indeed, light nicotine cigarette smokers (who smoke comparable amounts to the average participant in the current study) do not show withdrawal effects when abstinent (<u>Shiffman, 1988</u>). Nevertheless, our conclusions rest on comparisons between the ON smoking session and OFF smoking session. Individuals will be in a relatively higher cholinergic state in the ON smoking session whether or not they experience withdrawal symptoms for the OFF session.

### Stimuli

The stimuli were 3D rendered rooms, each of which had a unique shape, different pieces of furniture, and a single painting **(Figure 2)**. This stimulus set was previously used in Ruiz et al. (2020). Below, we describe how the rooms were created, how paintings were selected, and how rooms and paintings were combined.

**Figure 2.**
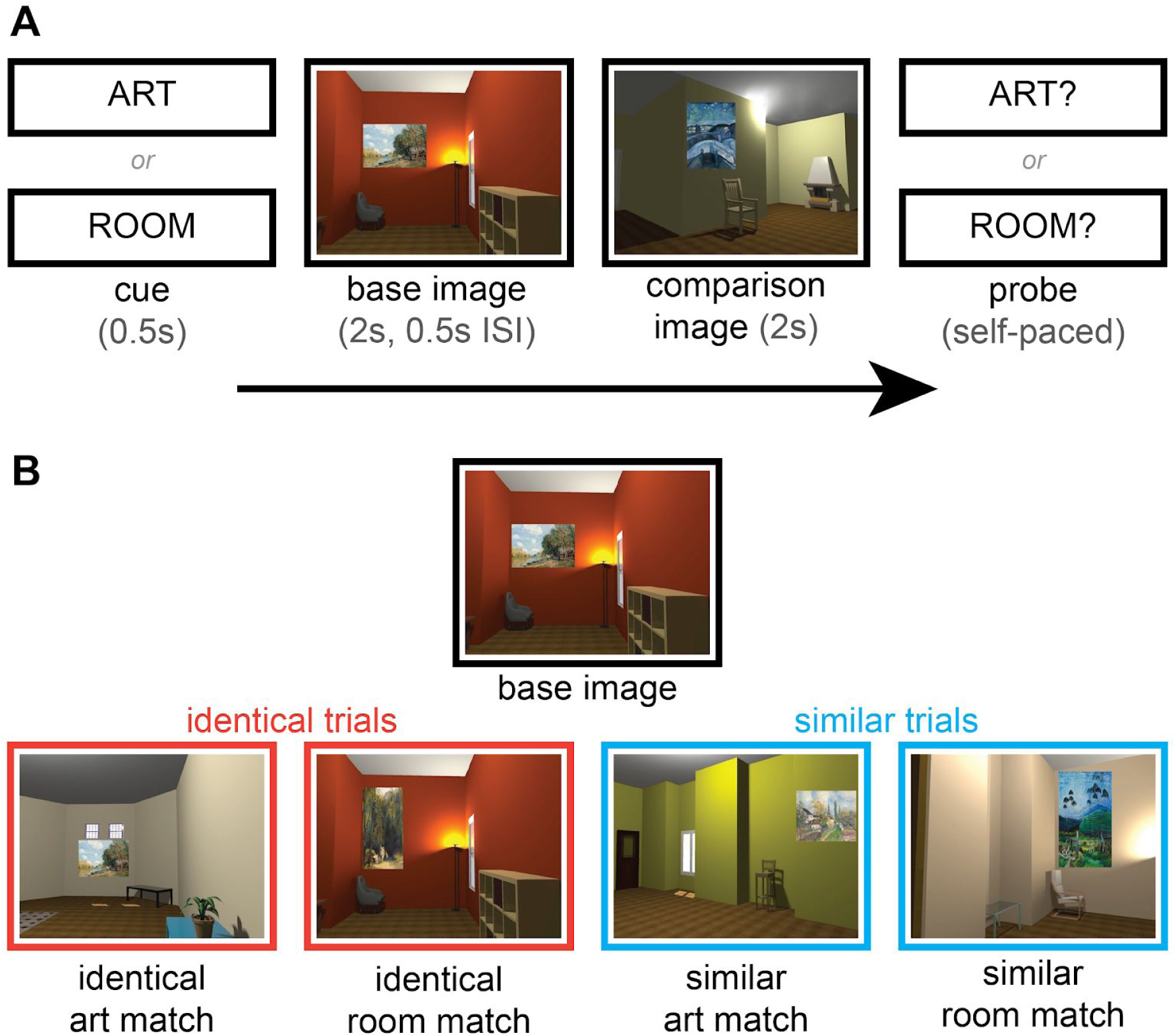
**(A)** Trial structure. On every trial, participants viewed two rooms, each containing one painting. Prior to trial onset, they were cued to attend to either the style of the paintings (ART) or to the layout of the rooms (ROOM). They then saw a base image and a comparison image. Finally, they received a probe (ART? or ROOM?). If the probe was “ART?”, participants had to judge whether the two paintings matched, i.e., whether they could have been painted by the same artist. If the probe was “ROOM?”, participants had to judge whether the two rooms matched, i.e., whether they had the same spatial layout. On valid trials, the initial cue matched the probe at the end of the trial. On invalid trials, the cue and the probe were different (e.g., “ART” cue and “ROOM?” probe, or vice versa). **(B)** Examples of art and room matches. An art match could either be a painting identical to that in the base image (identical art match) or a different painting that was painted by the same artist as the painting in the base image (similar art match). A room match could either be a room identical to that in the base image (identical room match) or a room with the same spatial layout as the base image, from a different perspective (similar room match). A non-matching image contained neither an art match nor a room match and could also be displayed as a comparison image, as in **(A)**. “Identical” trials involved presentation of a base image and one of the following: an identical art match, an identical room match, or a non-matching image. “Similar” trials involved presentation of a base image and one of the following: a similar art match, a similar room match, or a non-matching image. ISI = inter-stimulus interval.

Rooms were rendered using Sweet Home 3D (http://www.sweethome3d.com/). An initial stimulus set of 80 rooms was created. From these, a second version of each room was generated with a 30-degree viewpoint rotation (half clockwise, half counter-clockwise). These versions, which we refer to as “similar room matches”, were additionally altered so that some visual features changed while spatial geometry (i.e., wall angles, wall lengths, furniture layout) remained the same as the original image. Specifically, wall colors were changed and furniture pieces were replaced with a different exemplar of that furniture type (e.g., a chair was replaced with a different chair). Thus, each original image and its similar room match have the same spatial layout from a different perspective, but the images differ in their low-level visual features. An additional 10 rooms and their similar room matches were created for a practice run of the task.

A stimulus set of 160 paintings — 80 different artists with 2 paintings each — was selected from the Google Art Project (https://artsandculture.google.com/). The artists were chosen such that their styles were distinguishable, but paintings by different artists nevertheless had overlapping content (primarily outdoor scenes, some natural and some with man-made objects; some scenes also contained people). This ensured that participants could not use broad artistic categories (e.g., representational vs abstract art) as the basis for their assessments of style. The 2 paintings by each artist (a painting and its “similar art match”) were selected so that they were similar in terms of style, but with potentially different content. Thus, each painting and its similar art match have commonalities in the shades and variety of color, the level of detail, and the type of brushstroke, even if the content of the paintings differs. An additional 20 paintings were chosen from 10 artists (2 paintings each) for use in the practice run of the task. None of the stimuli used in the practice run appeared in the main experiment.

We then combined the paintings and rooms. Each of the 160 rooms for the main experiment (80 original rooms and their similar room match) were paired with 3 paintings, all by different artists. Likewise, each of the 160 paintings for the main experiment (80 original paintings and their similar art match) were paired with 3 different rooms, all with different spatial layouts. This generated a stimulus set of 480 unique images for the main experiment. For the practice run of the task, the 20 practice paintings and 20 practice rooms were combined using the same logic as above to create 60 unique images.

### Design

We followed the same experimental design as Ruiz et al., (2020). For the main experiment, 80 image groupings of 6 images each were created from the stimulus set of 480 unique images. There were 80 total trials in the main experiment, each of which used one image grouping. The groupings consisted of a “base image” (the first image to appear on a given trial), a “similar art match” (a room that contained a different painting by the same artist as that in the base image), a “similar room match” (a room with the same spatial layout as the room in the base image, viewed from a different perspective), an “identical art match” (an image with the identical painting that was in the base image), an “identical room match” (a room that was identical to that in the base image), and a non-matching image (an image with a painting made by a different artist and a room with a different spatial layout than the base image) **(Figure 2)**. Images containing an art match (either identical or similar) to the base image could not also be a room match (either identical or similar), and vice versa. An additional 10 image groupings were created using the same logic for the practice run of the task.

The 80 trials for the main experiment were divided into 40 “art” attention trials and 40 “room” attention trials. On “art” trials, participants were instructed to attend to the style of the paintings presented on that trial, and to assess their use of color, brushstroke, and amount of detail. On “room” trials, participants were instructed to attend to the arrangement of furniture and the wall angles of the rooms presented on that trial.

Within these attentional states, half of the trials were “identical” trials and half were “similar” trials. Identical trials involved presentation of a base image and either: an image with an identical painting (“identical art match”), an image with an identical room (“identical room match”), or a non-matching image. Similar trials involved presentation of a base image and either: an image containing a different painting by the same artist (“similar art match”), an image containing a room with the same spatial layout from a different perspective (“similar room match”), or a non-matching image.

Each trial consisted of an “ART” or “ROOM” cue presented for 0.5 s, followed by a base image for 2.0 s, an interstimulus interval for 0.5 s, the comparison image for 2.0 s, and finally a probe, “ART?” or “ROOM?”. The probe stayed on the screen until the participant responded **(Figure 2A)**. The cue instructed participants to attend to either the style of the paintings in the images (“ART”) or to the layout of the rooms in the images (“ROOM”). Potential comparison images included either an identical art match, an identical room match, a similar art match, a similar room match, or a non-matching image **(Figure 2B)**. Participants were to respond “yes” if there was a match in the probed dimension and “no” if there was not a match. Specifically, participants were to respond “yes” to an “ART?” probe if there was an identical art match *or* a similar art match present on that trial, and “no” otherwise. Participants were to respond “yes” to a “ROOM?” probe if there was an identical room match *or* a similar room match present on that trial, and “no” otherwise. Responses were made on the keyboard using the 1 key for “yes” and the 2 key for “no”. There was no specific instruction to respond as fast as possible.

To measure whether our attentional manipulation (i.e., “ART” vs “ROOM” cue) was successful, we included both valid trials and invalid trials in the experiment (Posner, 1980). On valid trials (80% of trials), the cue at the beginning of the trial matched the probe at the end (e.g., both “ART” or both “ROOM”). On the remaining 20% of trials (invalid trials), the probed dimension and the cued dimension did not match (e.g., participants were cued to attend to art and were probed about a room match, or vice versa). If our attentional manipulation is effective, performance should be better on valid vs. invalid trials (Posner, 1980).

On each trial, the second (comparison) image could be one of the following: (1) a cued match (i.e., an art match [either identical or similar] on a trial with an art cue; a room match [either identical or similar] on a trial with a room cue); (2) a non-cued match (i.e., an art match [either identical or similar] on a trial with a room cue; a room match [either identical or similar] on a trial with an art cue); or (3) a non-matching image (neither an art match nor a room match).

On valid trials, the cued (and probed) match was presented on 50% of the trials, the non-cued (and non-probed) match was presented on 25% of the trials, and a non-matching image was presented for the remaining 25% of the trials. For invalid trials, the probed (but not cued) match was presented on 50% of the trials, the cued (but not probed) match was presented on 25% of the trials, and a non-matching image was presented on the remaining 25% of the trials. Therefore, across all trials the correct answer was “yes” half the time and “no” half the time.

The main experiment was presented in 8 blocks of 10 trials each, where all trials within a block had the same attentional cue (“ART” or “ROOM”). Attentional cues alternated across blocks (4 “ART” blocks and 4 “ROOM” blocks; order counterbalanced across participants). Similar and identical trials were intermixed throughout the experiment. Intermixing these trial types ensured that participants had to attend to the entire painting or room layout on each trial and attempt to extract multidimensional information, because they did not know in advance whether a potential match would be identical or similar. Thus, differences in performance across these trial types could not be a result of different attentional strategies during the presentation of the base image.

Participants received feedback at the end of each block. This feedback depicted the percentage of correct responses and a qualitative assessment based on performance level (e.g., “Wow! You are doing amazingly well! Keep it up!”, “You are doing very well! Keep it up!”, “You are doing ok! Keep it up!”, “This task is challenging, but keep trying!”).

The practice run of the experiment consisted of 10 trials. 5 of these trials began with an “ART” cue and 5 with a “ROOM” cue. Of these 10 practice trials, half were identical trials and half were similar trials. 8 of the trials were valid and 2 were invalid. Participants received feedback after every 5 trials (i.e., after the “ART” block and after the “ROOM” block).

### Procedure

The study took place over two sessions, which were scheduled roughly 7 days apart based on the participant’s availability (*M* = 8.16 days, *SD* = 3.58; range = 3-25 days). Participants were instructed to smoke at least 1 cigarette within 1 hour prior to their “ON” session and abstain from smoking for at least 12 hours prior to their “OFF” session **(Figure 3)**.

**Figure 3.**
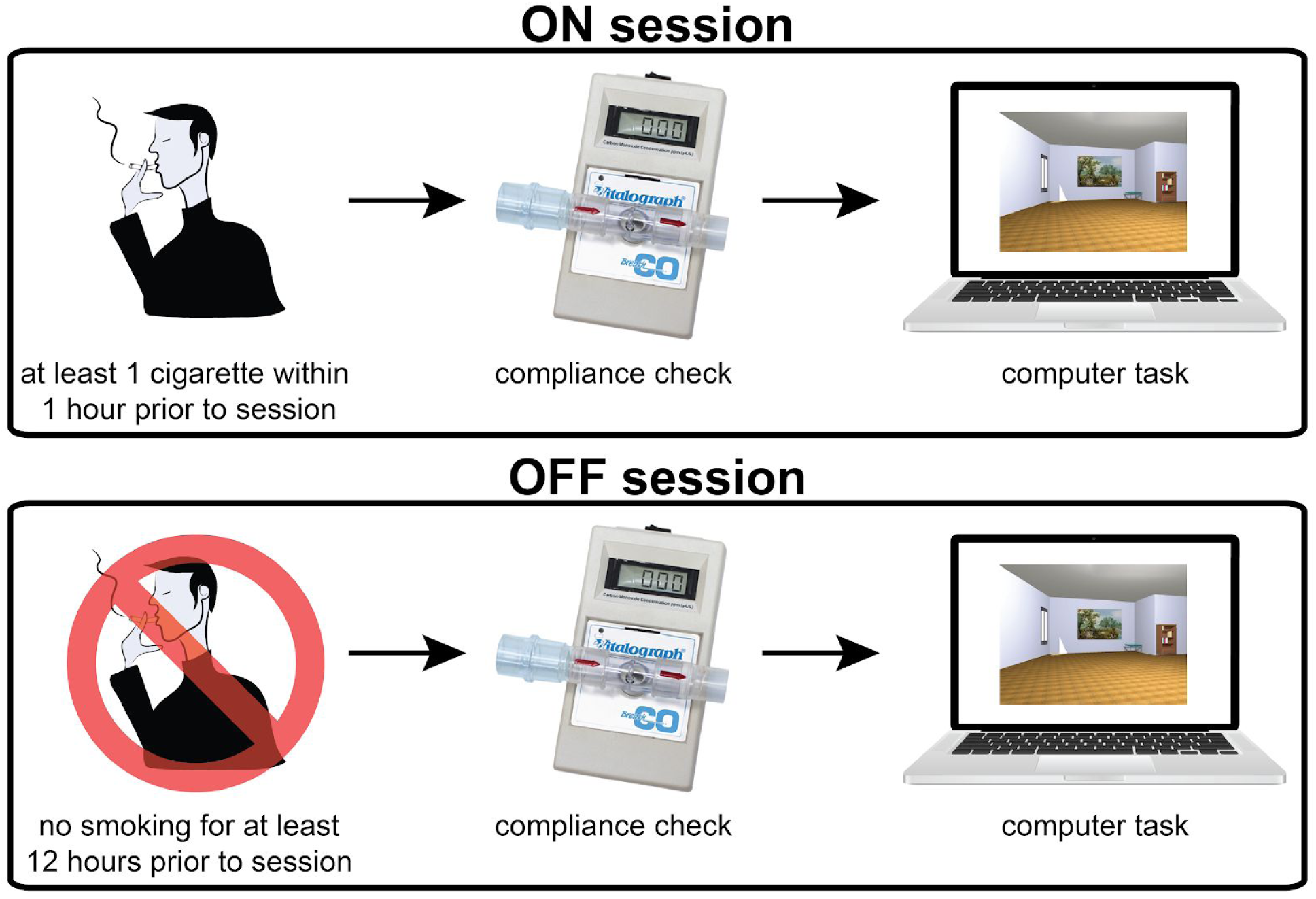
Study procedure. The study consisted of two sessions. For the “ON” session, participants were instructed to smoke at least 1 cigarette within 1 hour of the session’s start time. For the “OFF” session, participants were instructed to abstain from smoking for at least 12 hours prior to the session’s start time. Session order was counterbalanced across participants. Upon arrival for both sessions, participants were tested for compliance via an assessment of expired breath carbon monoxide (CO) with a breath CO monitor. Participants then received instructions for the task, were shown sample images of potential match types, completed a practice run, and completed the main experiment.

Nicotine reaches the brain within 10-20s of inhaling from a cigarette. Blood nicotine concentrations peak within minutes and then slowly decline, with a half-life of approximately 2 hours (Benowitz, Hukkanen, & Jacob, 2010; Berridge et al., 2010). The task used in the current study takes approximately 30 minutes to complete (after 10 minutes to obtain consent and provide instructions). Thus, the ON session was completed within a time period for which blood nicotine levels should be high, and within the half-life of nicotine

The order of “ON” and “OFF” sessions was counterbalanced across participants. The same stimulus set was used for both experimental sessions, but the assignment of stimuli to attentional task (art vs. room), trial type (identical vs. similar), and trial validity (valid vs. invalid) was randomized for each session.

Upon arrival for their first session, participants gave written informed consent, filled out a demographics form, and completed the Fagerstrom Test for Nicotine Dependence. A subset of participants (n = 23) completed additional neuropsychological examinations for a separate study; these test results are not reported here. At the beginning of both sessions, participants filled out a short questionnaire inquiring how many cigarettes they had smoked in the last hour and the last 12 hours. For the ON session, participants reported smoking 1-3 cigarettes within the last hour (*M* = 1.21, *SD* = 0.46) and 1-12 cigarettes within the last 12 hours (*M* = 2.36, *SD* = 1.88). For the OFF session, all participants reported smoking 0 cigarettes within the hour prior to the session. One participant reported smoking 1 cigarette exactly 12 hours prior to the session; the remaining participants reported smoking 0 cigarettes in the past 12 hours.

Participants were then tested for compliance via an assessment of expired breath carbon monoxide (CO) with a Vitalograph Breath CO monitor (Vitalograph, Lenexa, KS). This measurement produces a readout of CO level in parts per million (ppm). Expired breath CO provides a quick and non-invasive indirect measure of tobacco smoke intake (Jarvis et al., 1980; Wald et al., 1981) and can be used as an index of nicotine levels (Jarvik et al., 2000; Russell et al, 1978; Vollstädt-Klein et al., 2011; Vossel, et al., 2011). Our use of expired breath CO as a proxy for nicotine levels is supported by the relatively high correlation observed between these variables in several studies. For example, Vossel et al. (2011) report a correlation of 0.94 between blood cotinine (a nicotine metabolite) and expired breath CO. Second, Jarvik et al. (2000) report correlations of 0.83 – 0.98 between nicotine levels in the blood and expired breath CO. Correlations between expired breath CO and carboxyhemoglobin levels in the blood are also quite high, around 0.98 (Jarvis et al., 1980; Wald et al., 1981). Higher carboxyhemoglobin levels are also linked to higher nicotine levels (Russell et al., 1978). Together, these findings suggest that expired breath CO can be used as a fairly good indicator of nicotine levels.

Because there is debate regarding the particular CO ppm level that is consistent with smoking vs abstaining (Cropsey et al., 2014; Ernst et al., 2001; Perkins, Karelitz, & Jao, 2013; Yamamoto et al., 2013), we used a simple criterion that CO ppm must be higher for the ON vs OFF session for a participant to be included in the study (ON session ppm: *M* = 8.14, *SD* = 4.63; OFF session ppm: *M* = 3.06, *SD* = 1.97). Using this criterion ensures that participants smoked more prior to their ON session vs their OFF session, but we cannot be certain that participants completely abstained from smoking prior to their OFF session. The comparison of ON vs OFF sessions can be considered a comparison of more vs less smoking without any change to our conclusions. It is worth noting, however, that the average OFF session ppm (3.06) is close to the suggested “abstinent” cut-off of 3 based on large-scale studies of nicotine cigarette smokers (Cropsey et al., 2014, Javors, Hatch, & Lamb, 2005).

Furthermore, our conclusions do not depend on an assumption that nicotine smokers were in a “normal” cholinergic state in the OFF session and an “above-normal” cholinergic state in the ON session. It could be that nicotine smokers experience some withdrawal in the OFF session, and are therefore in a “below-normal” cholinergic state in the OFF session and a “normal” cholinergic state in the ON session. We consider this unlikely given that no participants reported moderate or high nicotine dependence, and light cigarette smokers (who smoke comparable amounts to the average participant in the current study) do not show withdrawal symptoms when abstinent (<u>Shiffman, 1988</u>). Nevertheless, we cannot rule out that some individuals experienced withdrawal. In either case, however, participants were in a higher cholinergic state in the ON vs OFF session, and that is the only requirement for testing our hypotheses.

After the CO assessment, participants received instructions for the attention task and were shown examples of a base image and all the comparison image types (identical art match, identical room match, similar art match, similar room match). Participants were not given particular strategies to use, and multiple different approaches have been reported (e.g., for the room task, participants may choose to focus on wall angles, furniture layouts, or both). After viewing sample images, participants then completed a practice run of 10 trials. Participants were required to achieve 80% accuracy or higher to proceed to the main experiment. All participants met this requirement. Participants then completed the main experiment. Both the practice run and the main experiment were conducted via the Psychophysics Toolbox 3 in Matlab (http://psychtoolbox.org/). The same procedure was followed for both sessions (i.e., the example images were shown, and practice run completed, for both sessions; for the second session, this served as a refresher). Upon completion of the second session, participants were debriefed and compensated.

### Statistical Methods

#### General Approach

We used Bayesian generalized mixed models to determine which task variables affect behavioral performance. These models were implemented via MCMC sampling in the Stan language using the rstanarm package in R (rstanarm v2.19.2, Goodrich, Gabry, Ali, & Brilleman, 2019; rstan v2.19.2, Stan Development Team, 2019; R v3.5.0, R Core Team, 2018). We opted to use a generalized logistic mixed model rather than a traditional analysis of variance (ANOVA) because only the former model can capture uncertainty in participant-level estimates of binary yes-no performance. Because ANOVA requires a continuous outcome variable, such an analysis would require each participant’s performance to be summarized in terms of a continuous measure (e.g., d’, A’, percent correct, or similar). But this approach ignores the sampling error of these summary statistics. For example, A’ calculated from 16 trials is more certain than A’ calculated from 4 trials. A generalized logistic mixed model allows us to fit binary (yes/no) trialwise responses in our attention task, thus enabling us to adjust for uncertainty in our estimates of A’ according to the number of trials in the task design. The results of logistic mixed models (beta estimates for each independent variable of interest) can be interpreted similarly to F-values from an ANOVA, where larger magnitude statistics imply a larger effect. Furthermore, we report 95% credible intervals (CIs) for all main and interaction effects on estimated A’, which can be interpreted similarly to p-values. 95% CIs that do not include 0 reflect differences between conditions that are statistically significant at p < .05. For completeness, we also report a traditional ANOVA in Supplemental Materials.

Unless otherwise specified, the mixed models were run with the following rstanarm settings: 4 sampling chains, with initial parameter guesses drawn from [-1, 1]. Each chain ran for 2000 total iterations, with the first 1000 iterations designated for warm-up and the second 1000 iterations for sampling. Unless otherwise specified, we report point estimates for each coefficient at the mean value of the posterior distribution across all sampling iterations and chains, and two-tailed 95% credible intervals of the same posterior distribution. All models demonstrated sufficient mixing of chains, fewer than 10 post-warmup divergent transitions for any single parameter, and an effective N of at least 10% of the sampling iterations for every parameter, diagnosed visually using the shinystan package (v2.5.0, Gabry, 2018).

#### Task Performance: Trialwise Match Detection and A’

We modeled task performance as a Signal Detection Theory (SDT) process using multilevel logistic regression. Our specific model makes the assumption that each participant’s latent “evidence strength” distributions for match present and match absent trials are overlapping logistic distributions, each with the same scale parameter, which indexes the spread of each distribution. This is analogous to two normal distributions with the same standard deviation, but with heavier tails than normal distributions. The model yields estimates of SDT discriminability and bias parameters by modeling the probability that, for a given trial, a participant endorses a match being present for match-absent and match-present trials respectively (DeCarlo, 1998; Rouder & Lu, 2005). Our multilevel trial-wise model of participant responses contrasts with single-level individual difference analyses on point estimates of signal detection metrics calculated from raw performance data. It has the advantage of (a) using the individual trial-level data to estimate SDT parameters at the group level, i.e., for the average participant, (b) accurately characterizing individual-difference heterogeneity in those parameters, and (c) potentially relating these to covariates (see <u>Bolger, Zee, Rossignac-Milon & Hassin, 2019</u>). The model directly estimates the locations of participants’ underlying “signal” and “noise” distributions, and it allows traditional summary measures of behavioral sensitivity to be extracted from the model’s estimates and interpreted as traditional signal detection metrics. As such, we report model-estimated sensitivity, operationalized as A’ (Donaldson, 1992; Ruiz et al., 2020), in conjunction with untransformed coefficient estimates from the model. A’ is a measure of behavioral sensitivity for which 0.5 indicates chance performance and 1.0 indicates perfect performance.

We used a logistic link function to model P(match endorsement = “yes”) as a function of the other predictors. A logistic link function is appropriate in this context because the ROC distribution implied by A’ is consistent with underlying logistic “signal” and “noise” distributions when performance is above chance but below ceiling (Macmillan & Creelman, 1996). As such, A’ estimated from a logistic regression model should have the same properties as A’ calculated from raw performance data and can be interpreted equivalently.

We included main effects of all experimental conditions **(Table 1, model 1)**, as well as fixed effect terms for all possible two-way and three-way interactions. Including these terms allowed model-estimated performance, indexed by the parameter estimate for match status, to vary as a function of different combinations of experimental conditions, without overspecifying the model. Similarly, we included a random intercept, as well as random effects for all experimental conditions and two-way interactions. We did not include any further random interaction effects to avoid overspecifying the model. All predictors were effect-coded, with each predictor’s two levels coded as -0.5 and +0.5. Effect-coding allows for ANOVA-like interpretation of model parameters, such that the intercept can be interpreted as a grand mean and the main effects are estimated at the mean of each of the other predictors. We set weakly informative Cauchy priors with mean = 0 and scale = 2.5 for all terms. Cauchy priors are well-suited for the coefficients of Bayesian logistic regressions, as they provide the regularizing benefits of a bell-shaped prior while allowing large values of coefficients to be estimated when appropriate, e.g. when responses are separated (Gelman, Jakulin, Grazia, Pittau, & Su, 2008).

**Table 1.** Overview of the main analysis approaches. Model #1 and Model #5 were the analyses of primary interest (results from which are shown in Figures 4, 5, and 6). The remaining models tested secondary hypotheses. In particular, considering the A’ results along with the results of the response time (RT) models (#2 and #4) allowed us to determine if there were speed/accuracy trade-offs, but we did not have strong hypotheses about how nicotine might affect RTs in this task. The models incorporating trial validity (#3 and #4) allowed us to ensure that the attention manipulation was effective (faster and more accurate responses on valid vs. invalid trials). In Model #1, the main effect of match status reflects the magnitude of behavioral sensitivity across all experimental conditions. Differences in sensitivity between experimental conditions are indexed by interaction terms between match status and the other experimental conditions (i.e., attentional state, condition, and smoking session). The same logic holds for Model #3, with the added experimental condition of trial validity. In Model #2, differences in response time (RT) for hits, correct rejections, misses, and false alarms are indexed by main effects of match endorsement, match status, and the interaction of match endorsement and match status. RT differences between experimental conditions, conditional on response accuracy, are indexed by interaction terms between match endorsement, match status, and the other experimental conditions (i.e., attentional state, condition, and smoking session). The same logic holds for Model #4, with the added experimental condition of trial validity. For Model #5, A’ OFF is re-centered at 0.5, so that the regression intercept is estimated when A’ OFF smoking is at chance performance. Additionally, for Model #5, results hold when A’ OFF is not included as an independent variable.

| Model | Model Type | Dependent Variable | Independent Variables | Interaction Terms | Variable Coding | Description |
| --- | --- | --- | --- | --- | --- | --- |
| 1 | trialwise multi-level model | endorse_match<br>0 = no<br>1 = yes | match_status<br>(absent, present)<br>attentional_state<br>(art, room)<br>condition<br>(identical, similar)<br>smoking_session<br>(OFF, ON) | all main effects<br>(fixed and random)<br>all two-way interactions<br>(fixed and random)<br>all three-way interactions<br>(fixed) | effect coding<br>(-0.5 for first level of variable, 0.5 for second level) | primary model to assess A' across the experimental manipulations |
| 2 | trialwise multi-level model | logRT<br>(log-msec) | endorse_match<br>(no, yes)<br>match_status<br>(absent, present)<br>attentional_state<br>(art, room)<br>condition<br>(identical, similar)<br>smoking_session<br>(OFF, ON) | all main effects<br>(fixed and random)<br>all two-way interactions<br>(fixed and random)<br>all three-way interactions<br>(fixed and random) | effect coding<br>(-0.5 for first level of variable, 0.5 for second level) | similar to model #1, but for response times |
| 3 | trialwise multi-level model | endorse_match<br>0 = no<br>1 = yes | validity<br>(invalid, valid)<br>match_status<br>(absent, present)<br>attentional_state<br>(art, room)<br>condition<br>(identical, similar)<br>smoking_session<br>(OFF, ON) | all main effects<br>(fixed and random)<br>all two-way interactions<br>(fixed)<br>all three-way interactions<br>(fixed) | dummy coding for validity<br>(0=invalid, 1=valid)<br>effect coding for everything else<br>(-0.5 for first level of variable, 0.5 for second level) | similar to model #1, but to assess whether A' differs for valid vs invalid trials |
| 4 | trialwise multi-level model | logRT<br>(log-msec) | validity<br>(invalid, valid)<br>endorse_match<br>(no, yes)<br>match_status<br>(absent, present)<br>attentional_state<br>(art, room)<br>condition<br>(identical, similar)<br>smoking_session<br>(OFF, ON) | all main effects<br>(fixed and random)<br>all two-way interactions<br>(fixed)<br>all three-way interactions<br>(fixed) | dummy coding for validity<br>(0=invalid, 1=valid)<br>effect coding for everything else<br>(-0.5 for first level of variable, 0.5 for second level) | similar to model #2, but to assess whether response times differ for valid vs invalid trials |
| 5 | between-participants<br>linear<br>regression | A' ON - A' OFF | CO ppm ON - CO<br>ppm OFF<br>A' OFF | all main effects | CO ppm ON - CO<br>ppm OFF not<br>re-centered or<br>scaled<br>A' OFF re-centered<br>at 0.5 | examine whether<br>individual differences<br>in nicotine smoking<br>predict behavior |

In order to generate more directly interpretable test statistics from our model, i.e. in units of signal detection performance as opposed to inverse logit units, we used rstanarm’s posterior_linpred() function to extract inverse-logit-transformed posterior estimates of P(match endorsement = “yes”) at every level of every predictor from each iteration of the posterior distribution. By treating the posterior fixed-effect estimate of P(match endorsement = “yes” | match status = present) as the posterior-estimated hit rate, and P(match endorsement = “yes” | match status = absent) as the posterior-estimated false alarm rate, we could then use these fixed-effect hit and false alarm rates to calculate group-level posterior estimates of A’, as well as A’ differences between various conditions of interest.

To assess model fit and validate subsequent inferences from the model, we calculated the errors between mean posterior predicted A’ and raw A’ for each participant in each condition. We examined median absolute error, as it does not up-weight extremely large errors, across participants in each task condition. Median absolute error was below 0.07 A’ units across participants in all task conditions.

#### Task Performance: Response Times

We modeled log-transformed response times with a multilevel linear regression. For this regression, we used the same base model formula as we did for the match detection (i.e., A’) model, but with an additional term for whether or not the participant endorsed a match being present on that trial **(Table 1, model 2)**.

For the same reasons as we effect-coded other predictors, match endorsement was effect-coded in the response time model to allow coefficients to be estimated at the grand mean of “match present” responses and “match absent” responses. We also accounted for differences in accuracy in the response time model by including two- and three-way interactions with the match endorsement and match status predictors, for both fixed and random effects, allowing the model to estimate differential effects for correct responses (hits, correct rejections) and incorrect responses (misses, false alarms). We set weakly informative normal priors with mean = 0 and standard deviation = 10 z-units for the intercept term, and standard deviation = 2.5 z-units for all other terms.

While the coefficients for our response time model are directly interpretable in log-RT units, for maximal interpretability we exponentiated each posterior estimate of log-RT extracted from posterior_linpred() to yield posterior distributions of estimated response time in milliseconds in each task condition, and calculated test statistics on RT differences between conditions.

#### Task Performance: Valid vs. Invalid Trials

In order to verify that participants were indeed following cue instructions and shifting their attention to the art features or room features of each trial, we ran two additional models predicting match detection/A’ and response time as a function of probe validity (valid vs. invalid). In these models, we also included main effect predictors of all other task conditions **(Table 1, models 3 & 4)**.

We dummy-coded probe validity with invalid trials set to 0 and valid trials set to 1. Because our primary models included only valid trials, we coded probe validity as 0-1 so that the main effect coefficients for our validity models would be reported for invalid trials, as those data are not characterized in the primary models.

We included main effect terms of the other task conditions so that main effects of probe validity would be estimated at the grand mean of those other conditions. We also included all possible two- and three-way fixed effect interactions between predictors, allowing us to model differential effects of probe validity on A’/RT in different experimental conditions. We included random effects only for the intercept and main effect terms, to allow those estimates to vary between participants, without over-specifying the model. All other model estimation parameters were identical between the two match detection models and the two response time models respectively.

#### Supplementary trialwise models

Our primary model was set up in a factorial manner, such that each task sub-condition fell into 1 of 8 bins: 2 [condition: identical or similar] x 2 [attentional state: art or room] x 2 [smoking session: OFF or ON]. Such a design assumes that some consistent construct underlies performance for each of the sub-conditions that share a factorial level, e.g., some consistent process underlies performance on all similar match trials relative to all identical match trials. However, any model specification carries with it particular side effects, which may be advantages or limitations depending on the context. In this case, by using interaction terms to match our model to the factorial design of the task, estimates are assumed to be more similar for factorial conditions sharing a level. For example, the estimates for similar art trials OFF smoking are regularized to be closer to estimates for other similar trials, other art trials, and other OFF smoking trials. Such a model allows us to make true factorial inferences, arguably the most powerful, about the effects of our manipulations.

However, the constraining effects of a factorial model specification may cause the model to *underestimate* the magnitude of a simple effect. For example, if there were a true underlying on-smoking improvement in performance, but only for similar room trials, the absence of the simple effect in the other three trial types (i.e., identical art, identical room, similar art) would cause the model to estimate the magnitude of the simple effect for similar room trials too conservatively, thus reducing our ability to detect such an effect.

An alternative model specification that mitigates this concern is a dummy-coded model. In a dummy-coded model, each combination of predictor levels is instead treated as an independent level of a single variable. For example, instead of treating condition (identical vs. similar) and attention (art vs. room) as two interacting predictors, a dummy-coded model might have one predictor with four levels: identical-art, identical-room, similar-art, and similar-room. As such, simple effects are less constrained by the model to be similar for overlapping conditions. For this reason, we estimated a supplementary model with two predictors: one predictor for trialwise match status (present vs. absent), and one predictor for task condition with 8 levels, one each for the OFF and ON smoking sessions for identical art, identical room, similar art, and similar room trials **(Table 2, model 1)**, to probe for simple effects on A’ separately for each sub-condition.

**Table 2.** Overview of supplementary analysis approaches. These models were run to ensure that the results we observed in our main analysis approaches (Table 1) were robust. Supplementary models #4 and #5 are reported in Supplemental Materials.

| Model | Model Type | Dependent Variable | Independent Variables | Interaction Terms | Variable Coding | Description |
| --- | --- | --- | --- | --- | --- | --- |
| **Supp 1** | trialwise multi-level model | endorse_match<br>0 = no<br>1 = yes | match_status<br>(absent, present)<br>task_subcondition<br>(identical art ON, identical art OFF, identical room ON, identical room OFF, similar art ON, similar art OFF, similar room ON, similar room OFF) | all main effects<br>(fixed and random)<br>all two-way interactions<br>(fixed and random) | effect coding for match_status<br>(-0.5=absent, 0.5=present)<br>dummy coding for task_subcondition<br>(7 binary predictors:<br>predictor 1: 1=similar room OFF, 0=all others<br>predictor 2: 1=similar art ON, 0=all others<br>predictor 3: 1=similar art OFF, 0=all others etc.<br>NB: similar room ON is 0 for all predictors) | supplementary model similar to main model #1, but with dummy coding for the 8 sub-conditions of interest |
| **Supp 2a-d** | trialwise multi-level models | endorse_match<br>0 = no<br>1 = yes<br>(separate models for identical art, identical room, similar art, similar room) | match_status<br>(absent, present)<br>smoking_session<br>(OFF, ON) | all main effects<br>(fixed and random)<br>all two-way interactions<br>(fixed and random) | effect coding<br>(-0.5 for first level of variable, 0.5 for second level) | supplementary model similar to main model #1, but with separate models for each attentional state x condition combination |
| **Supp 3** | between-participants linear regression | A' ON - A' OFF | CO ppm ON - CO ppm OFF | all main effects | CO ppm ON - CO ppm OFF not re-centered or scaled | supplementary model similar to main model #5, but does not include a regressor for A' OFF |
| **Supp 4** | between-participants linear regression | A' ON - A' OFF | CO ppm ON - CO ppm OFF<br>A' OFF<br>number of years the individual has been smoking | all main effects | CO ppm ON - CO ppm OFF not re-centered or scaled<br>A' OFF re-centered at 0.5<br>years smoking not re-centered or scaled | supplementary model similar to main model #5, but includes a covariate for chronic nicotine use |
| **Supp 5** | repeated measures analysis of variance | A' | attentional_state<br>(art, room)<br>condition<br>(identical, similar)<br>smoking_session<br>(OFF, ON) | all main effects<br>all two-way interactions<br>all three-way interactions | N/A | analysis of A' as a function of the experimental manipulations, as a supplementary comparison to multi-level models |

To further reduce model constraints, we estimated another set of four supplementary models, each with match status (present vs. absent) and smoking session (off vs. on) as predictors **(Table 2, models 2a-d)**. One model each was estimated for identical art, identical room, similar art, and similar room trials respectively. These models are the most liberal in allowing the estimated effect of smoking session on performance to differ across attentional states and conditions.

#### Individual Differences

In addition to our primary trialwise model of task performance treating smoking session as a binary predictor, we conducted between-participants linear regressions to investigate possible dose-dependent effects of smoking nicotine on detection performance.

For each individual, we first obtained their CO ppm difference for the ON vs OFF smoking session. This difference score provides a measure of how recently or how much they smoked for their ON session, correcting for “baseline” ppm in the OFF session. We also calculated each individual’s raw A’ difference between the ON and OFF smoking sessions, separately for each of the four tasks: identical art, identical room, similar art, and similar room. Thus, higher values indicate greater performance enhancement on nicotine. Finally, for each task, we regressed A’ difference as a function of CO ppm difference across individuals **(Table 1, model 5)**. These regressions included a nuisance regressor of each participant’s “baseline” A’ in the OFF session, to adjust for possible regression to the mean between sessions. In particular, using A’ in the OFF session as a nuisance regressor allows us to examine smoking-related improvements in A’, while adjusting for the fact that low-performing individuals have more chance to improve than high-performing individuals. However, our results are very similar when A’ in the OFF session was not used as a nuisance regressor (**Table 2, model 3;** for discussion of alternative approaches, see Allison, 1990; <u>Castro-Schilo & Grimm, 2017</u>; Cronbach & Furby, 1970; Montgomery, Nyhan, & Torres, 2018).

We report coefficient estimates and associated partial Pearson correlations for the effect of CO ppm difference on A’ difference from each task’s regression, as well as differences in partial correlations between tasks. If nicotine smoking enhances attentional performance in a given task, then the individuals who show the greatest CO ppm difference should show the greatest performance enhancements in the ON nicotine session.

To allow our standard error and confidence interval estimates from these regressions to account for the hierarchical nature of our data, even when using a non-hierarchical model, we bootstrapped sampling distributions for the coefficient estimates of the regressions using hierarchical nonparametric bootstrap resampling (Carpenter, Goldstein, & Rasbash, 2003; for a neuroscience application see Saravanan, Berman, & Sober, 2020). For each of 500 bootstrap iterations, we first randomly drew participant IDs with replacement, to achieve a total N equal to that of our original participant group. Then, for each of those participant IDs, we resampled with replacement trial outcomes within each factorial task level (match status, condition, attentional state, smoking session) from that participant’s raw data, to achieve a trial count within each condition equal to that of the original task design. We then broke up our resampled datasets into four sub-datasets by condition x attentional state (identical art, identical room, similar art, similar room) and re-ran the regressions of interest for each resampled sub-dataset. For each resampled regression, we extracted the coefficient and partial Pearson correlation for the CO ppm predictor. Finally, we constructed bootstrap sampling distributions of these extracted coefficients and correlations by aggregating their values over all resampled datasets. We report 95th percentile confidence intervals from each coefficient/correlation’s bootstrapped sampling distribution. To construct bootstrap distributions of the differences in partial correlations between tasks, e.g., similar room correlation - similar art correlation, or similar room correlation - identical room correlation, we took the difference between the pair of partial correlation values within each resampled dataset. As with the bootstrap distributions of individual coefficients and partial correlations, we aggregated over all resamples to yield a bootstrap sampling distribution of 500 difference values for each pair of correlations.

Our hierarchical bootstrapping method, where participant IDs are resampled first, and trialwise responses are resampled second, functionally reduces the number of comparisons in the above analyses. Because the same “participants” make up all the data for a given bootstrap iteration, the bootstrapped statistics for each task are not totally independent. Over all iterations, this produces non-independent sampling distributions for the effect of CO ppm on performance within each task. This reduces the effective number of comparisons, and thus the risk of false discovery due to chance (see Derringer, 2018).

Finally, to examine whether chronic nicotine use might affect our results, we conducted analyses that incorporated self-reported years of smoking as an additional nuisance regressor. The effects of CO ppm ON-OFF on A’ ON-OFF hold in this new analysis (**Table 2, model 4;** for further discussion, see Nicotine Use Covariates in Supplemental Materials).

## Data Availability

Stimuli, experiment code, analysis code, and data can be found at https://github.com/alylab/artmuseNicotine.

## Results

### Valid vs. Invalid Trials

We first examined whether our attentional manipulation was successful by comparing performance on valid vs. invalid trials. If attention is effectively engaged by the cue at the beginning of the trial (i.e., “ART” or “ROOM”), then performance should be more accurate and faster when the probe matches the cue (i.e., valid trials) than when it does not (i.e., invalid trials; Posner, 1980). To that end, we used a Bayesian mixed model to examine performance across experimental conditions as a function of trial validity (valid vs. invalid trials). These analyses confirmed that participants were more accurate on valid vs. invalid trials, as indexed by a main effect of validity (beta = 2.39, 95% CI [2.10, 2.69]; estimated A’ difference = 0.173, 95% CI [0.0836, 0.298]). We also found an interaction of validity and condition on accuracy, such that participants showed a smaller valid vs. invalid boost in A’ for identical trials relative to similar trials (beta = -1.45, 95% CI [-2.03, -0.874]; identical trials: estimated A’ difference = 0.132, 95% CI [0.0762, 0.202]; similar trials: estimated A’ difference = 0.214, 95% CI [0.127, 0.320]). Furthermore, an analysis of log-transformed response times (RTs) confirmed that participants were faster on valid vs. invalid trials (beta = -0.371, 95% CI [-0.441, -0.301]; estimated RT difference = -413 msec, 95% CI [-750, -94.9]).

Taken together, these results suggest that attention was effectively engaged by the cue at the beginning of the trial: Participants were more accurate and faster on valid vs. invalid trials. Having verified that participants modulated their attention based on trial instructions, we next focused analyses on valid trials.

### Art vs. Room and Identical vs. Similar Trials

We next examined how individuals performed across conditions (identical vs. similar trials) and attentional states (art vs. room) in Bayesian mixed models predicting performance as a function of identical vs. similar condition, art vs. room attention type, and on vs. off smoking session (see **Figure 4** for model outputs). As expected, A’ was substantially higher on identical vs. similar trials (**Figure 5**), as indexed by a main effect of condition (beta = -2.83, 95% CI [-3.27, -2.40]; estimated A’ difference = 0.0962, 95% CI [0.0373, 0.158]). We did not find a comparable main effect of condition on log-RTs (beta = 0.0196, 95% CI [-0.045, 0.0846]; estimated RT difference = 11.9 msec, 95% CI [-263, 204]).

**Figure 4.**
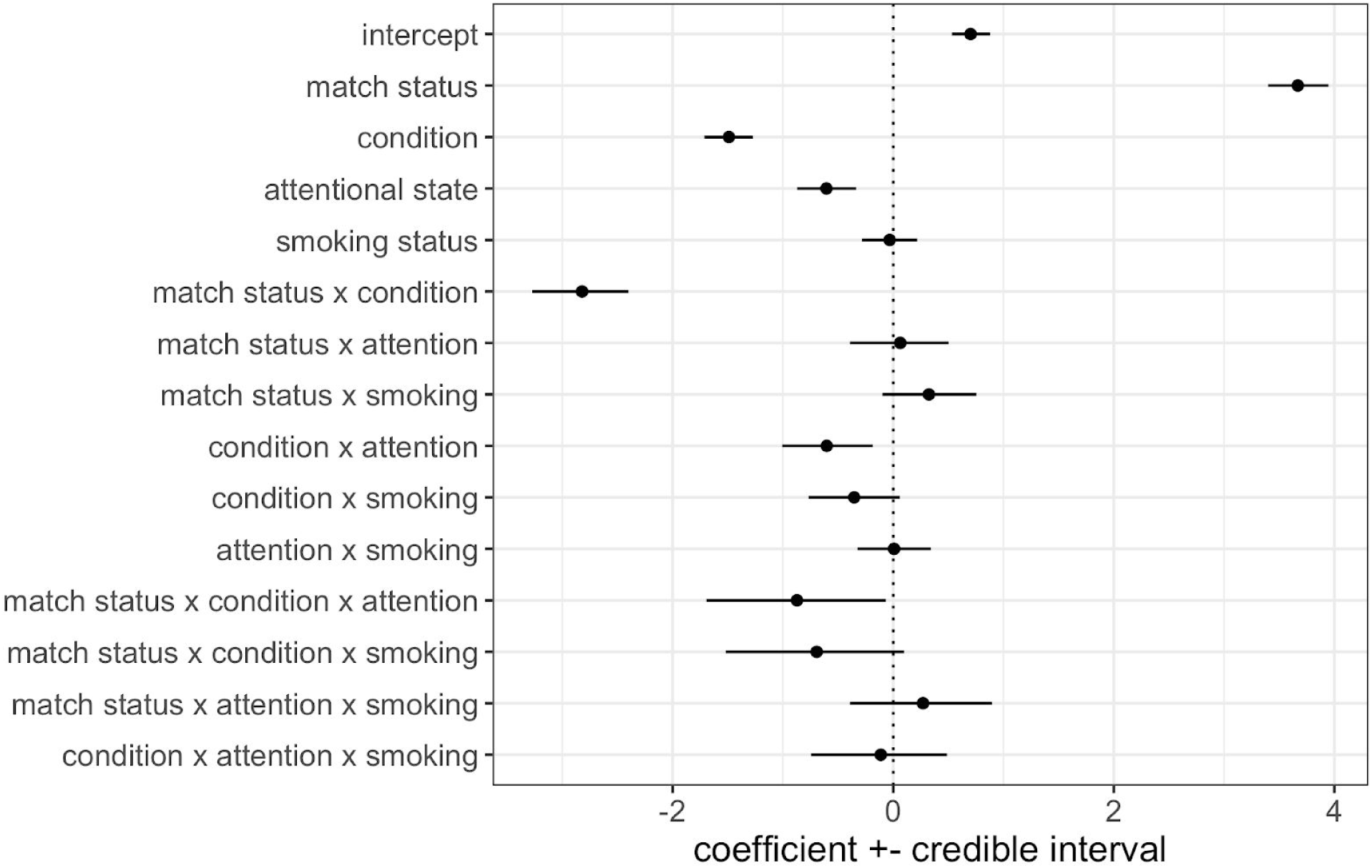
Coefficient estimates of the model for trialwise match detection performance (i.e., A’). In this model, the main effect of match status reflects the magnitude of behavioral sensitivity across all experimental conditions. Differences in sensitivity between experimental conditions are indexed by interaction terms between match status and the other experimental conditions (i.e., attentional state, condition, and smoking session). Accordingly, predicted A’ (shown below in **Figure 5**) was generated from these coefficients by treating the posterior estimate of P(match endorsement = “yes” | match status = present) as the hit rate, and P(match endorsement = “yes” | match status = absent) as the false alarm rate, in the A’ formula. Error bars = 95% credible intervals.

**Figure 5.**
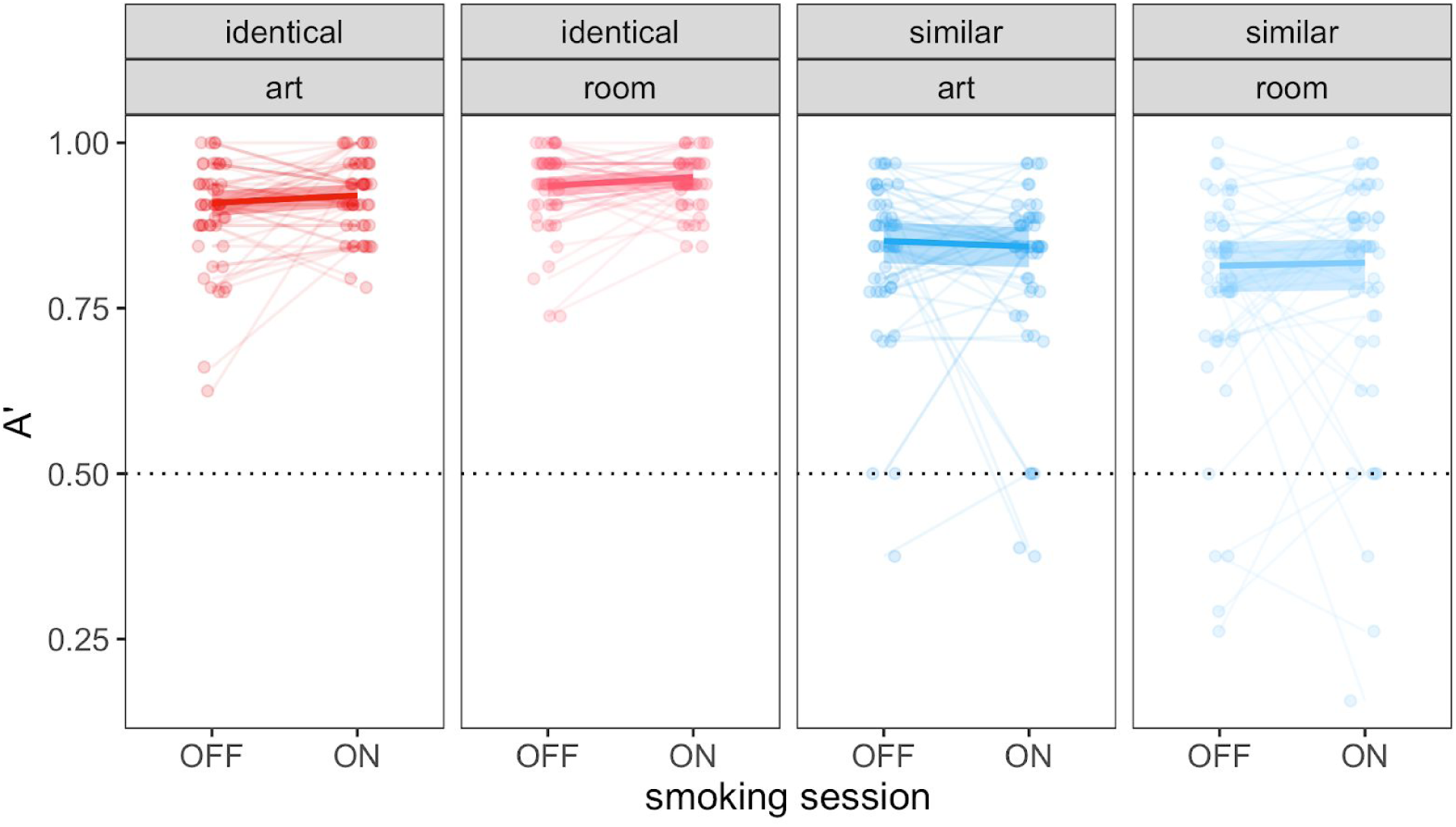
Behavioral performance (A’). Participants’ performance did not differ between OFF and ON smoking sessions for any trial type (each panel = one trial type). Points and faint lines indicate raw A’ values for each participant. Heavy lines and error ribbons indicate group-level model-predicted A’ ± 95% credible interval. Dashed horizontal line indicates chance performance (A’ = 0.5).

In contrast, performance was not meaningfully different between art and room attentional states, neither for A’ (beta = 0.0623, 95% CI [-0.393, 0.502]; estimated A’ difference = -0.00215, 95% CI [-0.0692, 0.0436]) nor for log-RTs (beta = -0.066, 95% CI [-0.162, 0.0324]; estimated RT difference = 38.2 msec, 95% CI [-304, 291]).

However, for A’, there was an interaction between condition and attentional state (**Figure 5**), beta = -0.875, 95% CI [-1.69, -0.0662]. A’ on identical trials was higher for room vs. art attention (estimated A’ difference = 0.0266, 95% CI [0.00697, 0.0466]), but A’ on similar trials did not meaningfully differ between room vs. art attention (estimated A’ difference = -0.0309, 95% CI [-0.0767, 0.0123]). There was no such interaction for log-RTs, (beta = 0.0201, 95% CI [-0.0584, 0.0933]).

### Comparison of ON vs OFF smoking sessions

We then turned to how performance on these tasks might be modulated by nicotine (**Figure 5**). We did not find a main effect of smoking session on A’, (beta = 0.325, 95% CI [-0.0985, 0.754]; estimated A’ difference = 0.00525, 95% CI [-0.0342, 0.0355]), suggesting that across trial types, A’ was not consistently higher during the ON session than the OFF session. We also did not find a main effect of smoking session on log-RTs (beta = 0.00293, 95% CI [-0.0701, 0.0713]; estimated RT difference = 2.79 msec, 95% CI [-191, 189]).

Next, we interrogated task-dependent effects of nicotine on performance. Because we initially hypothesized that nicotine would preferentially improve performance on similar room trials, we also examined two-way interactions between smoking session and condition (identical vs. similar) and smoking session and attentional state (art vs. room) on performance. We did not find a two-way interaction effect of smoking session and condition on A’ (beta = -0.699, 95% CI [-1.52, 0.0985]) or log-RT (beta = -0.00853, 95% CI [-0.0832, 0.0632]), nor did we find a two-way interaction effect of smoking session and attentional state on A’ (beta = 0.266, 95% CI [-0.393, 0.895]) or log-RT (beta = -0.0238, 95% CI [-0.0972, 0.0508]). Finally, an exploratory analysis examining hits and false alarms separately (rather than combined in an A’ score) also failed to show any differences as a function of ON vs OFF smoking sessions.

### Supplementary Models of Performance

The above analyses model the task as a 2 (condition: identical or similar) x 2 (attentional state: art or room) x 2 (smoking session: OFF or ON) factorial design. Although this has many advantages, there are also statistical disadvantages (see **Methods**). To assess whether our results vary based on statistical modeling approach, we also examined performance ON vs. OFF smoking with two other approaches. The first approach used a model with separate predictors for eight factorial levels of interest (i.e., the OFF and ON smoking sessions for identical art, identical room, similar art, and similar room trials). The second approach used four separate models to examine performance ON vs. OFF smoking in each task separately (one model each for identical art, identical room, similar art, similar room). We found that in these supplementary models, there was still no difference in performance between smoking sessions for any of the tasks. Thus, across multiple statistical modeling approaches, we failed to find a performance benefit ON vs OFF nicotine.

### Individual Differences

The above analyses treat smoking as a binary variable: they simply compared the ON and OFF smoking sessions. However, more precise information is available about individuals’ smoking, because for each session we obtained measures of expired breath carbon monoxide (CO) parts-per-million (ppm). This tells us how recently or how much an individual smoked, allowing us to investigate possible dose-dependent effects of nicotine smoking (e.g., Vollstädt-Klein et al., 2011; Vossel et al., 2011). If nicotine smoking enhances attentional performance, then the individuals who show the greatest CO ppm difference (in the ON vs. OFF smoking session) should show the greatest performance enhancements (in the ON vs. OFF smoking session). To investigate this, we ran four between-participants linear regressions predicting A’ difference as a function of CO ppm difference (ON vs. OFF), with one regression for each task: identical art, identical room, similar art, and similar room. The results below are not corrected for multiple comparisons, given that this was a planned analysis with a strong *a priori* hypothesis regarding the similar room task. Although we did not correct these results for multiple comparisons, our hierarchical bootstrapping approach (see **Methods**) effectively reduces the independence of the four comparisons, reducing the risk of false discovery.

Indeed, we found a modest positive correlation between these measures for the similar room task (beta = 0.0129 ppm units, bootstrapped 95% CI [0.000719, 0.0272]; partial Pearson correlation = 0.219, bootstrapped 95% CI [0.0126, 0.413]), such that a CO increase of 1 ppm from OFF to ON smoking predicted a 0.0129 unit increase in A’ from OFF to ON smoking (**Figure 6**). We obtained a very similar result when A’ in the OFF session was not included as a predictor variable (beta = 0.0151 ppm units, bootstrapped 95% CI [0.00139, 0.0333]; partial Pearson correlation = 0.216, bootstrapped 95% CI [0.0222, 0.417]). CO ppm differences did not predict performance enhancements on any other trial type (identical art: beta = -0.000942 ppm units, bootstrapped 95% CI [-0.00646, 0.00583], partial Pearson correlation = -0.0304, bootstrapped 95% CI [-0.295, 0.209]; identical room: beta = -0.00116 ppm units, bootstrapped 95% CI [-0.00677, 0.0034], partial Pearson correlation = -0.0841, bootstrapped 95% CI [-0.369, 0.225]; similar art: beta = -0.00102 ppm units, bootstrapped 95% CI [-0.015, 0.0111], partial Pearson correlation = -0.0259, bootstrapped 95% CI [-0.302, 0.203]).

**Figure 6.**
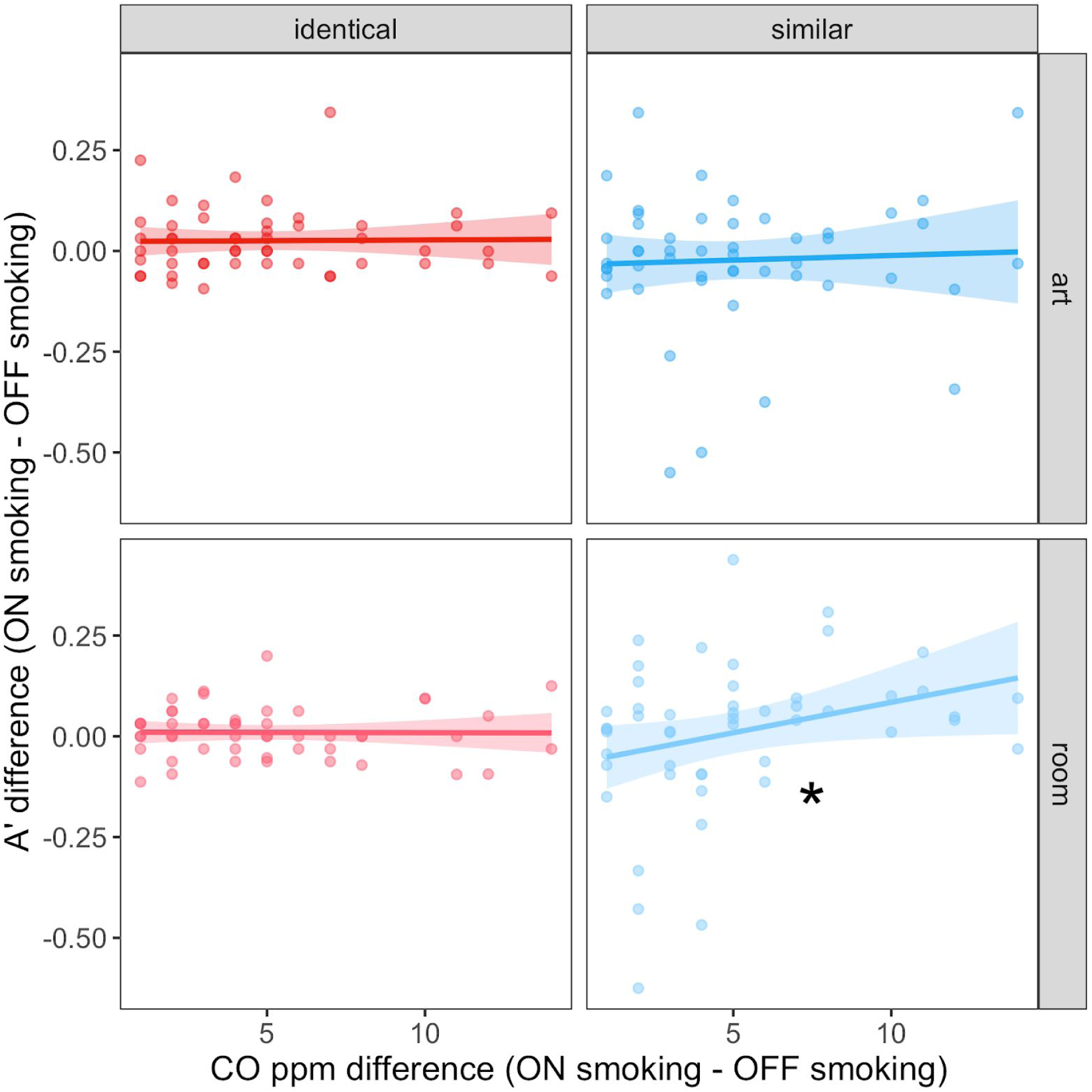
Dose-dependent effects of nicotine smoking on behavioral sensitivity (A’). On similar room trials (bottom right), participants who showed a greater increase in CO ppm from the OFF to the ON smoking session showed larger increases in A’ from the OFF to the ON smoking session. Participants did not show such an effect for identical art, identical room, or similar art trials. Within each panel, each point represents one participant; each participant appears in all four panels. * bootstrapped p < .05. Heavy lines and error ribbons indicate the line of best fit ± 95% confidence interval for ON-OFF A’ difference vs. ON-OFF CO ppm difference, displayed without adjusting for A’ OFF smoking for simplicity. Results are visually indistinguishable when adjusting for A’ OFF smoking.

Direct comparison showed that the partial Pearson correlation between CO ppm difference and A’ difference ON - OFF smoking for similar room trials was marginally higher than the partial Pearson correlations for the other three trial types (similar room > identical art: partial Pearson’s r difference = 0.249, bootstrapped 95% CI [-0.0702, 0.551]; similar room > identical room: partial Pearson’s r difference = 0.303, bootstrapped 95% CI [-0.0704, 0.701]; similar room > similar art: partial Pearson’s r difference = 0.245, bootstrapped 95% CI [-0.116, 0.604]). We note, however, that this is not strong evidence for a selective effect on similar room trials.

An exploratory analysis examining hits and false alarms separately (rather than combined in an A’ score) revealed that the smoking-related improvements on similar room trials were mostly driven by a reduction in false alarm rates (beta = -0.0153 ppm units, bootstrapped 95% CI [-0.0269, -0.00285]) rather than a change in hit rates (beta = 0.00374 ppm units, bootstrapped 95% CI [-0.0172, 0.028]). This result is broadly consistent with studies showing that nicotine can selectively reduce false alarms while not affecting hit rates (<u>Barr et al., 2007</u>; <u>Jubelt et al., 2014</u>), and nicotinic antagonists can increase false alarms with minimal effects on hits (Newhouse, Potter, Corwin, & Lenox, 1992). However, we made no *a priori* predictions about hit and false alarm rates, so this pattern of results should be replicated in future studies.

Finally, to examine whether chronic nicotine use might affect our results, we conducted analyses that incorporated self-reported years of smoking as an additional nuisance regressor. The effects of CO ppm ON-OFF on A’ ON-OFF hold in this new analysis (for further discussion, see Nicotine Use Covariates in Supplemental Materials).

## Discussion

Computational models of hippocampal function propose that cholinergic modulation prioritizes an externally oriented state in the hippocampus — a state that should promote attention to, and perception of, the outside world (Decker & Duncan, 2020; Hasselmo, 1995; Hasselmo, 2006; Hasselmo & McGaughy, 2004; Hasselmo, Wyble, & Wallenstein, 1996; Meeter et al., 2004; Newman et al., 2012). To test this, we had cigarette smokers perform an attention task (i.e., the similar room task) that recruits the hippocampus (Aly & Turk-Browne, 2016a; Aly & Turk-Browne, 2016b; Ruiz et al., 2020) while on nicotine and again while off nicotine. We also examined performance on three attention tasks that do not require an intact hippocampus (i.e., identical art trials, identical room trials, and similar art trials). We did not observe general improvement in performance on vs off nicotine on the hippocampally mediated similar room task, nor did we observe performance enhancements on the other trial types. However, the more an individual smoked nicotine cigarettes (as indexed by their expired breath carbon monoxide difference between the ON and OFF smoking sessions), the more they improved on similar room trials (as indexed by their A’ difference between the ON and OFF smoking sessions). This finding is broadly consistent with cholinergic modulation of externally oriented states in the hippocampus, and raises the possibility that only high levels of nicotine (or relatively large increases in cholinergic functioning) enhance hippocampally mediated attention and perception. Nevertheless, this effect was modest in size, and will be important to replicate in future work.

Why did analyses of individual differences in CO ppm reveal an effect of nicotine smoking on performance in the similar room task, while the comparison of ON vs OFF smoking sessions did not? Examination of individual differences on the similar room task (**Figure 6**) revealed that most individuals (34/50, more than would be expected by chance based on a sign test, *p* = 0.02) showed performance improvements on the similar room task in their ON vs OFF smoking session. However, a few individuals with low changes in CO ppm in the ON vs OFF sessions performed markedly *worse* in their ON session relative to their OFF session. This may have hurt our ability to detect an ON vs OFF change in behavior at a group level.

One caveat, however, is that we did not directly measure nicotine levels. Individuals who smoked more or smoked more recently should have higher nicotine levels, all else being equal — but cigarettes vary in nicotine content. Cigarettes with higher nicotine content are often associated with higher carbon monoxide levels (Benowitz, Jacob, Talcott, Hall, & Jones, 1986; Calafat et al., 2004; Lynch & Benowitz, 1987; Williams et al., 2007), and expired breath carbon monoxide has been used as an index of nicotine levels in studies of cigarette smokers (Vollstädt-Klein et al., 2011; Vossel et al., 2011). The use of expired breath carbon monoxide as a proxy for nicotine levels is further supported by findings of fairly high correlations (0.83 – 0.98; Jarvik et al., 2000; Vossel et al., 2011) between carbon monoxide levels and nicotine and/or cotinine levels in the blood (cotinine is a nicotine metabolite). Nevertheless, our findings should be replicated in future work that directly measures nicotine levels in order to provide stronger evidence.

Furthermore, the current work only manipulated nicotinic acetylcholine receptors, and muscarinic receptors are also important for balancing externally and internally oriented states in the hippocampus (Easton et al., 2012; Haam & Yakel, 2017; Hasselmo, 2006; Hasselmo & McGaughy, 2004; Newman et al., 2012). Thus, it is possible that stronger effects would have been observed if agonists for both types of receptors were used, or if acetylcholinesterase inhibitors were used to increase acetylcholine levels (Silver et al., 2008). Activation of both nicotinic and muscarinic receptors should prioritize externally oriented states in the hippocampus, although via distinct mechanisms. In particular, activation of nicotinic receptors enhances afferent input (e.g., responses to incoming sensory stimuli), and activation of muscarinic receptors suppresses excitatory feedback connections (e.g., those mediating memory retrieval; Hasselmo, 2006).

We did not observe cholinergic modulation of performance on attention tasks that do not require an intact hippocampus for accurate performance (i.e., identical art, identical room, and similar art trials; Ruiz et al., 2020). This null effect should not be over-interpreted. Nicotine acts throughout the brain and can modulate processing in visual cortex (Arroyo, Bennett, & Hestrin; 2014; Disney, Aoki, & Hawken, 2007; Hahn et al., 2009; Lawrence, Ross, & Stein, 2002). Thus, nicotine might have been expected to enhance performance on non-hippocampally-dependent tasks as well. It is possible that studies that administer higher amounts of nicotine, and/or manipulate muscarinic receptors as well, might observe effects where we did not. We discuss this in more detail below, where we highlight connections between our work and prior studies of cholinergic modulation of hippocampal function, visual cortex function, attention, and perception. We then consider additional limitations and future directions.

## Relation to Prior Work

Our work was inspired by studies that sought to test whether acetylcholine can toggle the hippocampus between externally and internally oriented states (Tarder-Stoll, Jayakumar, et al., 2020). For example, computational modeling work and research with non-human animals has found that high levels of acetylcholine prioritize attention / encoding states in the hippocampus, while low levels of acetylcholine prioritize memory retrieval (Easton et al., 2012; Hasselmo & Schnell, 1994; Hasselmo, 1995; Hasselmo & Barkai, 1995; Hasselmo et al., 1995; Hasselmo et al., 1996; Hasselmo & McGaughy, 2004; Meeter et al., 2004; Newman et al., 2012). Furthermore, pharmacological manipulations in humans have found that antagonists of muscarinic acetylcholine receptors impair new learning but do not affect recall of previously learned information (Atri et al., 2004). Finally, behavioral studies in humans have indirectly examined how acetylcholine might modulate the trade-off between encoding and retrieval by manipulating environmental novelty vs familiarity. Environmental novelty increases acetylcholine release (Giovannini et al., 2001), and should therefore promote hippocampal memory encoding. Conversely, lower levels of acetylcholine in familiar environments should promote hippocampal memory retrieval. Indeed, recent exposure to novel stimuli improves behaviors that depend on encoding precise memories (Duncan et al., 2012), while recent exposure to familiar stimuli improves behaviors that benefit from retrieval of past memories (Duncan et al., 2012; Duncan & Shohamy, 2016; <u>Patil & Duncan, 2018</u>).

However, to our knowledge, no research in humans has directly tested whether cholinergic agonists enhance hippocampally mediated attention and perception, in tasks with no demands on long-term memory. We aimed to do this by using an attention task that recruits the hippocampus in fMRI studies (Aly & Turk-Browne, 2016a; Aly & Turk-Browne, 2016b) and for which accurate performance depends on an intact hippocampus / medial temporal lobe (Ruiz et al., 2020). Contrary to our hypothesis, we did not observe general improvement on the hippocampally mediated similar room task when individuals were on vs off nicotine. Instead, we observed monotonic improvements in performance with more nicotine cigarette smoking, raising the possibility that higher amounts of nicotine ingestion (or relatively large increases in cholinergic functioning) might be needed to observe effects on hippocampal function. This finding is nevertheless generally consistent with computational models of acetylcholine modulation in the hippocampus.

One reason that we may not have observed robust effects of nicotine is that performance on the attention task used here might also benefit from some degree of internally oriented processing. That is, although attention/encoding generally requires more externally oriented processing than does memory retrieval, both internal and external modes likely contribute to some extent in both attention and memory tasks. In the current study, deciding whether the base image is an art or room match to the comparison image would benefit from maintenance of the base image in mind across the short inter-stimulus interval. This (very-short-term memory) might benefit from a hippocampal “retrieval” mode. Use of an attention task that requires less internally oriented maintenance, e.g., if the base and comparison images were simultaneously presented, may show larger performance benefits from a high cholinergic state.

Although our focus has been on cholinergic modulation of hippocampal function, many studies in humans have found that cholinergic modulation affects visual cortex function (for review, see Bentley, Driver, & Dolan, 2011) and improves attention and perception (for review, see Klinkenberg, Sambeth &, Blokland, 2011). For example, in visual cortex, nicotine modulates activity levels (Hahn et al., 2009; Lawrence et al., 2002) and acetylcholinesterase inhibitors reduce the spatial spread of responses (Silver et al., 2008). However, the effects of nicotine on visual cortex activity are not always consistent, with some studies showing increased (Lawrence et al., 2002) and some showing decreased (Hahn et al., 2009) activity. Moreover, nicotine cigarette smokers, relative to non-smokers, exhibit reductions of task-related activity in visual cortex during visual attention tasks (Vossel et al., 2011). Vossel et al., (2011) also found that nicotine cigarette smokers show increased parietal cortex activity, and faster response times to invalidly cued targets, as a function of expired breath carbon monoxide. This latter result suggests a potential role for nicotine in modulating attention via its effects on parietal cortex. Indeed, nicotine modulates the responses of parietal regions linked to attentional alerting and reorienting (Thiel, Zilles, & Fink, 2005).

Behaviorally, nicotine enhances attentional re-orienting (Thiel et al., 2005), reduces response times on stimulus detection and selective attention tasks (Ernst et al., 2001; Hahn et al., 2007; Hahn et al., 2009), and improves sustained visual attention (Mumenthaler et al., 2003). However, other studies have been unable to find behavioral attention improvements with nicotine (Giessing, Thiel, Rösler, & Fink, 2006; Griesar, Zajdel, & Oken, 2002; Impey, Chique-Alfonzo, Shah, Fisher, & Knott, 2013). Finally, nicotine can sometimes *hurt* performance: it can impair attentional selectivity, working memory accuracy, and visual scanning and attention (Heishman & Henningfield, 2000; Vangkilde, Bundesen, & Coull, 2011).

Thus, nicotine inconsistently modulates performance on attention tasks: it can help, hurt, or have no effect. A better understanding of the factors that lead to performance modulation vs not as a function of nicotine might shed light on why we observed no effects of nicotine on identical art trials, identical room trials, or similar art trials. One possibility is that only high levels of nicotine (or relatively large increases in cholinergic functioning) will enhance performance, consistent with our interpretation of the monotonic performance enhancement on similar room trials as a function of smoking recency / amount. This would also be consistent with studies that report a link between expired breath carbon monoxide levels and attentional biases (Vollstädt-Klein et al., 2011) and changes in brain activity during visual attention tasks (Vossel et al., 2011). Because we only tested individuals who were relatively light or moderate smokers, ingested nicotine levels might have been too low to see an effect on visual cortex function and concomitant behavioral improvements on identical trials and similar art trials.

Thus, one might speculate that the threshold for nicotine to affect visual cortex function may be higher than the threshold to see effects on hippocampal function. While there is no direct support for this, there is evidence that nicotine might differentially affect hippocampus vs visual cortex. For example, the effects of cholinergic modulation change across lower-order visual areas vs higher-order visual areas vs association cortex, in part because of differing expression profiles of nicotinic and muscarinic receptors, and differences in cholinergic innervation (Galvin, Arnsten, & Wang, 2018). Additionally, there are numerous subtypes of nicotinic acetylcholine receptors that vary in their sensitivity. Both high and low affinity nicotinic acetylcholine receptors are found in the hippocampus (Newman et al., 2012) and throughout the cortex (Arroyo et al., 2014). The density of receptors that are highly responsive to nicotine could affect the levels at which nicotine smoking starts to affect attentional functions in a given brain area. Given the variety of acetylcholine receptor subtypes and prevalence across different brain regions (Alkondon & Albuquerque, 2004), and the numerous ways in which cholinergic modulation may affect other neurotransmitters, it is not surprising that the mechanisms and roles of cholinergic function differ in hippocampus and sensory cortex (Metherate, 2004). Therefore, it is plausible that higher levels of nicotine are needed to affect attention processes in visual cortex vs hippocampus — but direct studies of this are required.

Another possibility for the null effects observed when comparing the ON vs OFF smoking sessions is that the tasks used here were not challenging enough to show robust performance benefits from nicotine. Using tasks with faster stimulus presentation times and/or more trials might have taxed individuals more, and allowed us to have more sensitivity to detect nicotine-related enhancements. A final possibility is that stronger manipulations of the cholinergic system are needed to show performance enhancements in the tasks used here. This could be accomplished by the use of acetylcholinesterase inhibitors, which increase synaptic acetylcholine levels and can thus enhance the action of both nicotinic and muscarinic receptors (e.g., Gratton et al., 2017; Kukolja et al., 2009; Rokem, Landau, Garg, Prinzmetal, & Silver, 2010).

## Limitations & Future Directions

Traditional pharmacological studies in humans — i.e., those using double-blind manipulations in healthy individuals — are logistically very challenging, and require careful medical prescreening, physiological measurements, and/or access to medical professionals to assist in the study (Heishman & Henningfield, 2000; Rokem et al., 2010; Wignall & de Wit, 2011). One major advantage of our approach of studying nicotine cigarette smokers is that studies that give nicotine to non-smokers have produced inconsistent effects on attentional behavior: some have found improvements (Thiel et al., 2005), some have found impairments (Heishman & Henningfield, 2000, Vangkilde et al., 2011), and some have found no effects (Giessing et al., 2006; Griesar et al., 2002; Impey et al., 2013; for review see, Newhouse, Potter, & Singh, 2004). This inconsistency may, at least in part, arise because of individual differences in responses to nicotine (Perkins, 1995). Additionally, administering nicotine to non- or never-smokers can produce dysphoria and other performance impairing effects (Foulds et al., 1997; Heishman & Henningfield 2000). Regular smokers show tolerance to these effects (Perkins et al., 1993). Thus, studying nicotine cigarette smokers who self-administer nicotine in a way that they are used to might help reduce the variable and sometimes disruptive responses to nicotine seen in non-smokers.

This is related to a second advantage of our approach, which is that the experiment was logistically very tractable and naturalistic: individuals were able to ingest nicotine as they usually would in their daily life. However, this comes with disadvantages that are not present in tightly controlled pharmacological studies. First, we did not control the amount of nicotine ingested. Individuals were allowed to smoke as much as they preferred prior to their ON session, and they could smoke whatever brand of cigarettes they preferred. Moreover, although individuals had to smoke within the hour prior to the ON session, individuals may have varied considerably in when they smoked within those 60 minutes. Thus, there was likely great variability in nicotine levels across individuals as they were doing our task — mirrored by the variability we observed in expired breath carbon monoxide levels. Future studies might find larger effects if they had individuals smoke the same type of cigarette at the same point in time prior to the session. Alternatively, future studies can allow these factors to vary, but take note of precise smoking times and cigarette brands, and use those as covariates in analyses. In this way, a middle-ground can be achieved between our naturalistic approach — which allowed smokers to follow their normal habits prior to the ON session — and the tightly-controlled pharmacological studies that are more commonly done.

Along those lines, another limitation of the current study is that it was not double blind. Both the participant and the experimenter knew which session was on nicotine and which was off nicotine. This concern is partly alleviated by performance measures that are objective (A’ and response times) rather than subjective. Furthermore, participants were not aware of our predictions — i.e., that we hypothesized an effect primarily for similar room trials. If nicotine improved performance on all trial types, it would not be clear if it was improving behavior directly via its neural effects or indirectly because of participant expectations of how they should perform on vs off nicotine. We did not observe such a general enhancement on all trial types and indeed, we did not observe any performance differences between the ON vs OFF sessions at a group level. This mitigates the concern that knowledge of nicotine ingestion affected our results. Future studies using this approach would benefit from having trial types for which researchers do not expect nicotine to improve performance, in order to better detect effects that may be driven by knowledge of smoking status.

Finally, future work that combines pharmacological approaches and fMRI will be important for shedding light on the neural mechanisms by which nicotine can enhance hippocampally mediated attention and perception. This is especially the case because some studies find no or weak behavioral effects of nicotine while observing robust neural differences (Giessing et al., 2006; <u>Thiel & Fink, 2008</u>; Thiel et al., 2005). Furthermore, combining fMRI and pharmacological approaches can provide insights into the mechanisms by which nicotine acts. The results reported here are broadly consistent with nicotine enhancing externally oriented states in the hippocampus, insofar as individuals who smoked more, or smoked more recently, showed the largest performance enhancements on similar room trials. But similar behavioral improvements might be expected if nicotine is enhancing visual cortex processing, with downstream consequences for hippocampal function. Thus, although the results are generally consistent with predictions of hippocampal computational models, other mechanisms can potentially give rise to the observed behavioral effects. Repeating this study in conjunction with fMRI can help clarify whether nicotine is enhancing spatial relational attention and perception via effects on the hippocampus, or via actions on visual cortex or other systems.

## Conclusion

We tested whether cholinergic modulation in humans enhances performance on attention and perception tasks that recruit the hippocampus. We find evidence that is generally consistent with cholinergic modulation of hippocampal attention and perception. Nicotine cigarette smokers who smoked more, or smoked more recently, showed greater performance enhancements on a hippocampally mediated attention task. More broadly, we suggest that nicotine cigarette smokers can provide a tractable approach for testing computational models of cholinergic functioning in the human hippocampus.

## Author Contributions

N.A.R. and M.A. designed research. N.A.R performed research. M.T. and N.A.R. analyzed data. N.A.R. and M.T. made figures. N.A.R, M.T., and M.A. wrote the paper.

We have no conflicts of interest to disclose.

This work was funded by an NSF CAREER Award (BCS-1844241) and a Brain & Behavior Research Foundation NARSAD Young Investigator Grant (27893) to M.A. We would like to thank Christopher Medina-Kirchner for valuable discussions about cholinergic manipulations, and Niall Bolger for valuable feedback about statistical methods. We are also grateful to Debby Song, Trevor Dines, and Allya Elgayar for assistance with data collection. Finally, we are grateful to the Alyssano lab meeting group for insightful feedback.

## Supplemental Materials

### Analysis of Variance

In addition to our analyses based on logistic mixed models, we conducted a more traditional repeated-measures analysis of variance (ANOVA) as a post-hoc, supplementary robustness check. This ANOVA is analogous to our main logistic mixed model (model #1 in **Table 1**), but uses participant-level A’ as the dependent variable, rather than trialwise yes/no responses. The independent variables were attentional state (art, room), condition (identical, similar), and smoking session (on, off), with all possible main effects, two-way, and three-way interactions. This ANOVA specification is roughly equivalent to a mixed model with only random intercepts for each participant.

Consistent with our logistic mixed model, the repeated-measures ANOVA revealed a main effect of condition (identical vs. similar; F(1, 391) = 100, p < .001), and a two-way interaction between condition and attentional state (identical/similar vs. art/room; F(1, 391) = 8.88, p = .0031). The remaining main effects and interactions were not meaningful (all F(1, 391) < 1.06, all p > .30). Any post-hoc analyses exploring the directionality of these effects are functionally equivalent to the post-hoc analyses on our logistic mixed models described in the main text. We thus do not report additional post-hoc analyses on this repeated-measures ANOVA.

### Nicotine Use Covariates

We collected a series of self-report measures of chronic nicotine use: the Fagerstrom Test for Nicotine Dependence (FTND), estimated cigarettes smoked/day, and years of smoking. To examine whether these chronic nicotine use variables might affect the relationship between CO ppm ON-OFF and A’ ON-OFF in the similar room task, we first conducted a principal components analysis (Supplemental Figure 1). This enabled us to explore the correlations between self-reported nicotine use measures and expired breath CO (OFF smoking, ON smoking, and the ON-OFF difference), which in turn allows us to identify covariates that contribute unique information to the between-participants regression. All variables were z-scored before submitting to principal components analysis using R’s stats::prcomp() function. The first two principal components together explained 79.3% of the total variance. FTND score, cigarettes/day, and all of the CO ppm measures contributed primarily to the first component, while years of smoking contributed to the second component only. The covariance between FTND score, cigarettes/day, and CO ppm makes intuitive sense, because individuals who smoke more cigarettes per day are likely to report higher levels of nicotine dependence and to smoke more prior to the ON session.

Given that FTND score and cigarettes/day covaried with CO ppm, while years smoking seemed to capture unique information, we opted to use only the years smoking variable as a covariate in the between-participants CO ppm model for similar room trials. This is well justified because using correlated predictors in regression can lead to coefficients that are uninterpretable (Farrar & Glauber, 1967). Adding years smoking as a covariate to the analyses revealed that reported years of smoking did not predict A’ ON-OFF (beta = 0.00136 ppm units, bootstrapped 95% CI [-0.0127, 0.0152]). Further, reported years of smoking did not fundamentally alter the effect of CO ppm on A’ ON-OFF (without covariate: beta = 0.0129 ppm units, bootstrapped 95% CI [0.000719, 0.0272]; with covariate: beta = 0.0139 ppm units, bootstrapped 95% CI [-0.00205, 0.0313]). Accordingly, the partial Pearson correlation between CO ppm difference and A’ difference ON - OFF smoking was very similar when including years smoking as a covariate (without covariate: partial Pearson correlation = 0.219, bootstrapped 95% CI [0.0126, 0.413]; with covariate: partial Pearson correlation = 0.224, bootstrapped 95% CI [-0.0391, 0.46]). We note that five participants were missing reported years of smoking, and were kept in the original regression, but excluded from the regression with the covariate. This reduction in sample size may contribute to the wider CIs observed for the effect of CO ppm in the covariate regression. This analysis therefore does not reveal strong evidence that prolonged nicotine use had an effect on our observed results.

**Supplemental Figure 1.**
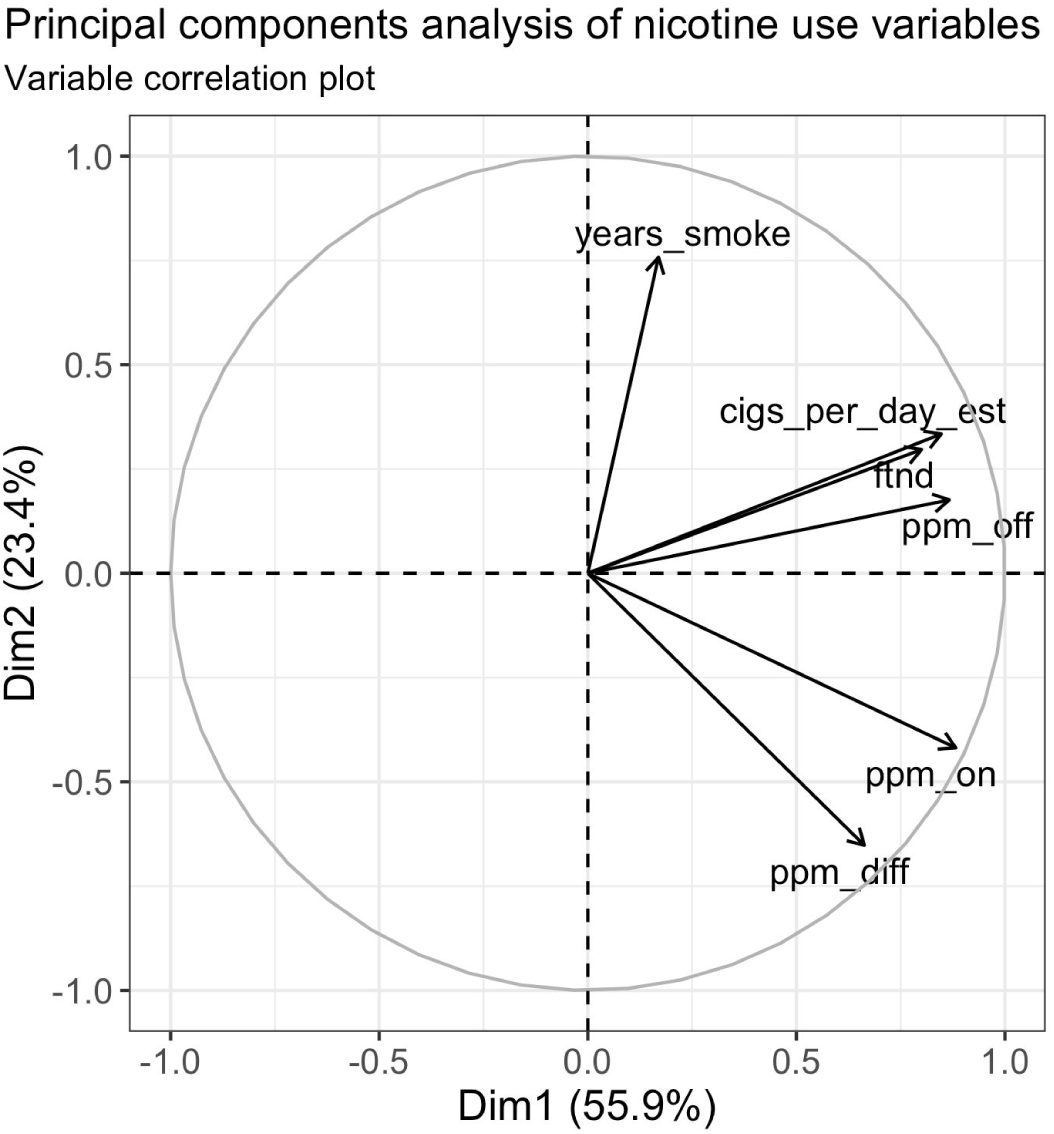
Principal components analysis on nicotine use measures. The first principal component explains 55.9% of total variance, while the second component explains 23.4%. Length of arrow indexes how well the first two principal components capture the variance in that variable. A variable whose arrow touches the unit circle is completely explained by some combination of the first two components, while a variable with a very short arrow is poorly explained by the first two components. Arrow angle indexes the ratio of contributions of the first two principal components to that variable. A variable whose arrow points along either the x or y axis selectively contributes to the first or second principal components respectively, while a variable whose arrow points at a 45-degree angle contributes equally to the first and second principal components. CO ppm OFF, cigarettes/day, and FTND score contribute almost exclusively to the first component, while CO ppm ON and CO ppm ON - OFF contribute a bit more to the first than the second component. Years of smoking contributes almost exclusively to the second component.

